# A sample-to-identification workflow to monitor animal and plant biodiversity using metabarcoding and real-time analysis

**DOI:** 10.64898/2026.10.07.756996

**Authors:** Lucas Esteban Wange, Javier Prado-Martinez, Pol Alentorn-Moron, Tomàs Marques-Bonet, David Juan, Esther Lizano

## Abstract

Environmental DNA (eDNA) has emerged as a transformative tool for biodiversity monitoring, offering a non-invasive means to detect and identify species across various ecosystems. However, despite its potential, current eDNA methodologies have significant limitations, including limited sensitivity, restriction to a specific taxonomic group, lengthy processing times, the need for specialized laboratory conditions and equipment, as well as computational bottlenecks in equipment, connectivity, and algorithms. Overcoming these limitations is crucial for the real-time monitoring of a high number of taxa in field-based scenarios. Here we present an integrated eDNA workflow from extraction to data analysis that mitigates several of these challenges through several technical innovations. Our approach adapts the Tagsteady protocol to Nanopore sequencing, integrates multiplex PCR for simultaneous plant and animal detection, and introduces RTBioScan, a real-time analysis pipeline employing hierarchical basecalling and OTU-level consensus generation to achieve high-accuracy taxonomic assignments during sequencing. Using a single set of universal primers, we integrate metabarcoding and barcoding into a cohesive protocol, utilizing universal markers for both animals and plants (cytochrome oxidase 1 region, COI, and Internal Transcribed Spacer 2, ITS2). This comprehensive approach can detect and identify a broad spectrum of species, from individual organisms to entire communities in a single workflow.

## Introduction

Biodiversity loss caused by human activities has been recognized as a major global crisis and has led to a 73% global decrease in vertebrate populations since 1970 (LPI 2024) (WWF 2024). To face this crisis, it is of utmost importance to have efficient, scalable tools to catalog and monitor species abundance in large areas over long periods of time. Although global efforts have started to tackle this enormous task, they are currently limited by the available technology. Environmental DNA (eDNA) has been proposed as a powerful tool to overcome these challenges (Baird and Hajibabaei 2012). Approaches like line transects, capture-release, bioacoustics, and camera traps have been critical in assessing biodiversity in the past and will remain a main source of information for biodiversity assessments; however, they each have their own set of intrinsic limitations in terms of scalability and sensitivity (Iacaruso et al. 2026). eDNA has the potential to complement these technologies and provide a more comprehensive picture of biodiversity (Baird and Hajibabaei 2012). Specifically, because it is universal and not limited to any specific taxonomic group, it can detect species beyond macroflora and macrofauna (Macher et al. 2026). This approach, also called Biomonitoring 2.0, is becoming more standardized and affordable and can provide invaluable information, but is still not widely adopted (Pawlowski et al. 2021). Particularly when looking at terrestrial biodiversity monitoring in more remote areas (Compson et al. 2020; Sahu et al. 2025; Newton et al. 2025). Here we present a rapid, sample-to-identification workflow to monitor animal and plant biodiversity using Nanopore real-time analysis.

Advances over the last decade have made eDNA the most promising emerging technology for biodiversity monitoring. Particularly, metabarcoding, which uses short DNA sequences to identify species in a complex sample by comparing the detected sequences to a database of labeled sequences, has been a crucial technology to infer species presence in a given environment (Compson et al. 2020). However, despite these technological improvements, some challenges remain (Pawlowski et al. 2021). In particular, when attempting large-scale monitoring in the most remote and biodiverse regions, implementation has been held back by a number of limitations, both experimental and computational (Sahu et al. 2025; Newton et al. 2025; von der Heyden 2023).

A typical eDNA metabarcoding experiment consists of the following broad steps: 1) sample collection and preservation, 2) DNA extraction, 3) barcode DNA enrichment by PCR, 4) sequencing library preparation, 5) DNA sequencing and data preprocessing, 6) species detection by searching against a database of reference sequences. For each of these steps, a multitude of solutions have been developed, and a thorough review is beyond the scope of this study (Bohmann et al. 2022, Hakimzadeh et al. 2024). However, we identified several key limitations. One such limitation is a relatively low sample throughput. This is partly due to the tedious DNA extraction protocols specific to each sample type (*e.g.*, water, soil, feces). The obtained DNA is limited in amount and may contain as little as one molecule of a given species. While high-throughput protocols for PCR amplification and library preparations that enable sample multiplexing through the use of DNA Tags, short endogenous sequences added to eDNA molecules, have been developed, they usually test for one barcode region and taxonomic group at a time (Buchner et al. 2021). A strategy to get the most out of the sample and test for several distant taxonomic groups is through the amplification of more than one target per reaction in a multiplex PCR (Cordier et al. 2019). This has the potential to be more economical with the source material and yield high specificity and sensitivity.

Primer design is a crucial determinant of specificity and sensitivity; while more targeted primers are very specific, they limit the scope of the biodiversity assessment; more general primers often suffer from nonspecific byproduct amplification (*e.g.*, bacterial COI or fungi in ITS2) (Hintikka et al. 2022).

Another limitation is that most protocols are optimized for Illumina sequencing. This impedes field analysis of eDNA and presents a hurdle for researchers in developing countries and remote areas where access to sequencing facilities is limited. A limitation that extends to the need for specialized equipment and a molecular biology laboratory (von der Heyden 2023). This is particularly important when considering that the global hubs of biodiversity tend to be located in the Global South, in more remote areas, far from the next research center (Sánchez Herrera et al. 2024).

Finally, long-read *in situ* sequencing using Nanopores has the potential to use longer barcodes. Most established DNA barcodes are short for two reasons: suspected DNA degradation and inherent limitations of Illumina sequencing (Bista and Lino 2026; Doorenspleet et al. 2025). However, DNA degradation has not been formally assessed in most environments, and longer DNA barcodes could potentially provide valuable additional information about populations.

In addition to these experimental limitations, we identified several computational challenges. Typically, the large amounts of data that would be collected in a large-scale biodiversity survey require access to high-performance computing resources (HPC) and the corresponding expertise to perform species detection and downstream analysis. As for the experimental limitations, these resources are unlikely to be available in remote areas with limited access to broadband internet connections and costly computational resources.

Lastly, species identification through eDNA requires a pre-existing database of DNA barcodes to identify taxa present in a given environment. Although global databases like BOLD (Ratnasingham et al. 2024) are continuously growing, the most biodiverse regions worldwide, such as rainforests, still have a high number of missing data. To close these gaps, barcoding efforts in these regions must be intensified. One strategy to increase database representation is to enable simple Nanopore-based barcoding (Hebert et al. 2025). Combining metabarcoding, the analysis of a complex eDNA sample, and barcoding, the *de novo* sequencing and identification of the DNA barcode from an individual specimen, in one experiment can be a valuable asset in this context (Erens et al. 2026).

With these limitations in mind, we aimed to generate a comprehensive workflow from DNA extraction to species identification in the most time- and cost-efficient manner while maintaining high sensitivity. To this end, a workflow for plant and animal biodiversity monitoring using eDNA should be 1) robust, 2) affordable, 3) portable and field-ready, 4) targeting a broad range of species, and 5) easy to use for non-experts.

Here, we aimed to detect both plant and animal species to obtain a global picture of Metazoa and Viridiplantae biodiversity in a single experiment. Based on a review of the recent literature and available databases, we targeted the mitochondrial cytochrome oxidase 1 region (COI) for the detection of animals and the nuclear ribosomal DNA barcode Internal Transcribed Spacer 2 (ITS2) for plants. These regions are ideal for comprehensive metabarcoding because they show sufficient variation to distinguish closely related species. We focused on ampliconic regions covering the best-studied regions of these loci, enabling us to discriminate a wide variety of species by taking advantage of the most comprehensive databases.

However, targeting metazoan mitochondrial barcodes poses several challenges in environmental DNA metabarcoding. The most common issue is the amplification of the bacterial COI gene, and consequently, the low proportion of on-target reads compared to bacterial-derived reads (Hintikka et al. 2022). To overcome this issue, we conducted a study to identify the most promising primers to increase specificity for animal DNA and avoid bacterial DNA amplification. Next, we systematically optimized these primers *in silico* to increase specificity while maintaining sensitivity.

To obtain the most information from a single, potentially limited, eDNA sample, we explored the use of Multiplex PCR by combining primers for ITS2 and COI in a single PCR reaction. We employed the previously published Tagsteady protocol, which marks amplicons with a unique combination of DNA Tags at the time of amplification, thus enabling the pooling of samples for the library preparation step (Carøe and Bohmann 2020). By using a ligation-based workflow, it reduces the risk of barcode swapping.

While the original Tagsteady protocol was developed for Illumina sequencing, we combine it with ligation-based Oxford Nanopore library preparation. Due to the portable nature of MinION sequencers, this protocol can be used outside a typical laboratory setting.

In addition, Nanopore sequencing has two additional features that can be useful in the context of eDNA metabarcoding. On the one hand, it overcomes the length restrictions of second-generation sequencing methods, allowing for longer DNA barcodes and thus more sequence information. On the other hand, Nanopore sequencing has the particularity that each pore sequences molecules sequentially, in contrast to the highly parallel sequencing-by-synthesis employed by Illumina and other second-generation short-read sequencing. This has the advantage that sequencing data are available as soon as a molecule has passed the Nanopore and thus can be analyzed while sequencing is still ongoing. While this feature has been mostly used to monitor sequencing throughput and quality decay, we exploited it to achieve species identification shortly after initiating the sequencing run, allowing us to monitor them in real time and hence be more efficient in the balance between read production and potential saturation in species detection. To this end, we developed RTBioScan, a real-time pipeline inspired by the round-based design of NanoRTax (Rodríguez-Pérez et al. 2022) but designed and implemented to support hierarchical basecalling, OTU-level consensus and multi-marker demultiplexing.

Here, we present a combined experimental and computational workflow able to deliver robust, accurate, sensitive, and fast eDNA-based sequence identification within hours. The experimental protocol can start from a variety of sample types and uses multiplex PCR and a streamlined workflow adapted to Nanopore-based sequencing (Figure 1A). The resulting libraries were then sequenced on a portable sequencing device, and the data were processed and analyzed in real time to produce highly accurate taxonomic identification using only a high-performance laptop computer (Figure 1 B).

**Figure 1.**
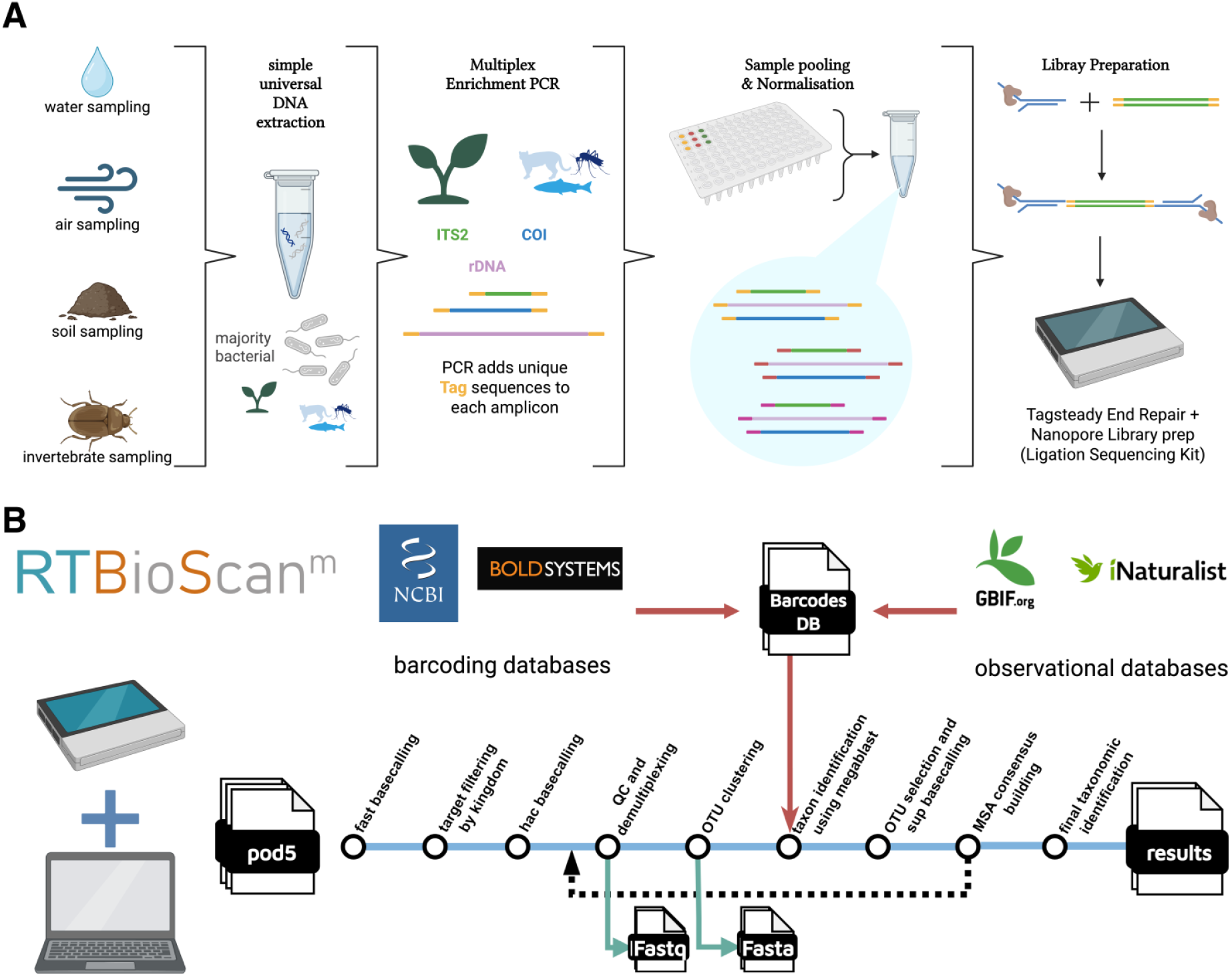
A unified workflow to monitor animal and plant biodiversity using environmental DNA metabarcoding. A) eDNA sample processing workflow. Bead-based eDNA extraction is used to obtain high-quality DNA from diverse sources. Next, enrichment PCR with 3’-tagged primers is performed to enrich animal and plant DNA. ITS2 for plants and COI for animals, were chosen and amplified simultaneously using multiplex PCR. A long ribosomal DNA region was targeted for long-read metabarcoding. Subsequently, the PCR products are pooled using a bead-based normalization strategy and end-repaired following the Tagsteady protocol. Ligation-based addition of Oxford Nanopore Sequencing adapters is performed to complete the library. The final pooled library is sequenced on MinION devices (Mk1B, Mk1C, and Mk1D) using R10.4 Flow Cells. B) Real-time data processing pipeline: RTBioScan. Our real-time data analysis pipeline takes the output of Nanopore sequencers as soon as it is generated. Sequencing output is processed in rounds of 200,000 reads. In each round, all reads are basecalled with the dorado fast model and screened with LAST against a kingdom-tagged marker database; only reads whose best hit is animal COI or plant ITS2 are retained, removing bacterial, fungal, and other off-target amplicons. Retained reads are basecalled a second time with the high accuracy (hac) model, length-filtered per marker, and demultiplexed by sample tag and primer with cutadapt. Reads are then clustered into operational taxonomic units (OTUs) with CD-HIT; OTUs are pre-classified by megablast against the reference barcode database, and up to 50 reads per OTU and sample are re-basecalled with the super accuracy (sup) model. A consensus barcode is built per OTU from these sup reads and classified by megablast. OTUs that remain stable across rounds are frozen, and their consensus sequences consolidated, so that later rounds only process new evidence.

## Materials and Methods

### Experimental methods

The selection of a method for extracting DNA from environmental samples was based on several considerations. First, it is easy to use and adaptable to field conditions, avoiding the need for reagents that have to be refrigerated and bulky equipment such as centrifuges. Second, it is effective in removing PCR inhibitors and is applicable to different types of environmental samples. Considering these requirements, we opted for a bead-based DNA extraction kit whose components do not require refrigeration except for the Proteinase K after resuspension.

### Sample collection

In this work, four types of sample collection were used. Soil samples were collected using a sterile spatula, air samples were collected passively with PES filters (0.22 µm and 0.45 µm pore sizes), water samples were collected passively with PES filters (0.22 µm and 0.45 µm pore sizes), and insect community samples were collected using yellow insect tape. Air and water samples were transferred to Zymo DNA/RNA Shield Lysis Tubes (Microbe) after exposure to the environment for 12 hours. Insect tape and soil were transferred to sterile containers and stored at -20 °C. For processing, 50 mg of soil was transferred to Zymo DNA/RNA Shield Lysis Tubes.

Soil sampling was performed in Collserola National Park, Barcelona, in December 2023. Air sampling was conducted in Barcelona, Nou Barris, on the balcony of a private apartment in May 2024. Water and insect sampling were performed in May 2024 in Mas Guimbau, Barcelona. Air, water, and insect samplers were exposed to the environment for 12-16 hours before collection.

### Environmental DNA extraction

Samples stored in DNA/RNA Shield Lysis Tubes (0.5 mm beads) were extracted using a magnetic bead-based isolation kit (Quick-DNA Magbead Plus Kit, Zymo Research) following the manufacturer’s protocol. Bead beating was performed for 2 min using a standard vortex mixer at maximum speed. In field-based scenarios, we also successfully performed the DNA extraction using a PortaLyzer, a self-made vortex mixer based on a battery-powered oscillating multitool (Peck et al. 2022). All samples were eluted in 50 μl of Elution Buffer (10 mM Tris, pH 8.5, 0.1 mM EDTA). Finally, the concentrations of all DNA extracts were measured with a Qubit fluorometer using a dsDNA High Sensitivity Assay Kit.

### Primer choice and optimization

The current metabarcoding literature was reviewed to identify primers with high specificity (low off-target) and high sensitivity (amplification of most species within the target taxon).

For plant metabarcoding, we focused on the UniPlant ITS2 primers (Moorhouse-Gann et al. 2018). It is designed to detect all Viridiplantae, and in comparison with the widely used chloroplast barcode rbcL, ITS2 shows higher interspecies sequence diversity and yields similar sensitivity and better specificity (G. S. Martin et al. 2024).

Animal metabarcoding has predominantly focused on mitochondrial DNA due to its higher copy number in animals. This focus has resulted in the generation of numerous mitochondrial barcode sequences for many animals. Especially for COI, as shown by the 19.9 million COI barcode sequences available in BOLD corresponding to 462,648 species (https://www.boldsystems.org, Snapshot 30th of June 2026). These extensive databases make COI, specifically its five-prime portion COI-5P, the most suitable target for broad universal animal detection in metabarcoding without focusing on specific taxa. Despite being the most extensively studied locus across animal species, there is substantial variability in the primer pairs and amplicons used to target it, many of which were originally optimized for barcoding. These primers often amplify different subregions of the locus, generating partially or non-overlapping amplicons that vary in length and sequence diversity, ultimately leading to markedly different sensitivities and specificities.

To improve the sensitivity and specificity of COI primers for selectively amplifying metazoan sequences while avoiding bacterial sequences, we started with a curated list of COI primer pairs previously tested in published studies. Among the two main COI barcoding regions (COI-5P and COI-3P), the COI-5P region has been more extensively characterized, with a larger number of species represented in sequence databases. Therefore, we focused on primer pairs targeting this region. To maximize sensitivity, we restricted our analysis to primer pairs that yield amplicons longer than 250 bp.

We evaluated each primer pair’s potential to recover metazoan versus bacterial sequences using the full set of COI entries available in the BOLD database. Specifically, we assessed the number of species containing exact matches (with no mismatches and accounting for primer degeneracy) to the forward and reverse primers, both independently and in combination. This distinction is critical: while single-primer matches indicate potential for partial recovery or off-target binding, only sequences matching both primers reliably indicate successful full-length amplicon recovery. Notably, longer amplicons are less likely to be fully represented in the database, but more likely to overlap at least partially with available sequences, which must be considered when interpreting sensitivity estimates.

An ideal primer pair should match a high number of metazoan species with both primers, while showing minimal or no amplification potential for bacterial sequences, particularly with both primers simultaneously. To systematically improve primer performance, we explored modified versions of each primer through a two-step optimization process.

First, to increase sensitivity, we sequentially introduced degeneracy at individual positions along the primer sequence. At each step, we retained modifications that increased metazoan recovery with a negligible impact on bacterial amplification (i.e., <1% increase). This process was repeated iteratively until no further gains were achieved without compromising specificity. Second, to enhance specificity, we performed the inverse operation by sequentially reducing degeneracy at each position and retaining modifications that improved specificity without significantly reducing sensitivity (again, using a <1% loss threshold). This iterative procedure allowed us to adapt previously published primer pairs—originally optimized for taxon-specific barcoding—to the broader demands of environmental metabarcoding using currently available COI data. We then experimentally compared the nine primer pairs with the highest *in silico* specificity and sensitivity to one of the most sensitive previously reported primer pairs BF3/BR2 (Elbrecht and Leese 2017). To this end, we processed a complex soil eDNA sample using the Tagsteady protocol with Nanopore sequencing. Primer pairs were compared using taxonomic yield per 1,000 reads and the proportion of reads assigned to Metazoa. These metrics quantify recovery and on-target enrichment in the tested sample.

All primers were appended with a unique 9-base-pair sequence tag, used to multiplex PCR reactions after amplification. Tag sequences were retrieved from Srivathsan et al. (Srivathsan et al. 2024). For each primer pair, we designed 16 uniquely tagged forward and reverse primers, enabling us to pool up to 256 PCR reactions in one run using combinatorial indexing.

### Multiplex PCR optimization and amplicon normalization

To minimize time and reagent costs and maximize the information per extracted DNA sample, we optimized a multiplex PCR assay to amplify the two target loci (COI and ITS2) in a single reaction. For this purpose, an initial gradient PCR to amplify each genetic locus individually was performed to determine the optimal annealing temperature for each primer pair. Reactions were carried out in a total volume of 25 μl using 1 μl of either Macaca silenus gDNA (17 ng/μl) for the COI primer pair or 1 μl of Sorghum bicolor (10.3 ng/μl) for the ITS2 primer pair. The reaction consisted of 12.5 μl of 2x QIAGEN Multiplex PCR Master Mix, 1.5 μl of each forward and reverse primer (10 μM each), the corresponding test DNA, and 8.5 μl of ultrapure water. Thermal cycling conditions included an initial denaturation step at 95 °C for 15 min, followed by 35 cycles at 95 °C for 30 s, with annealing temperature ranging from 44.5 to 64.5 °C for 90 s, and 72 °C for 90 s, and a final extension at 72 °C for 10 min. PCR products were visualized on an Agilent Bioanalyzer DNA 7500 Assay chip without further purification.

After determining a common optimal annealing temperature producing a single clear fragment for each primer set, we optimized the conditions to perform a multiplex PCR assay combining the two primer pairs. For this purpose, a mock sample (6X_DNA, 1 ng/μl) was prepared consisting of a mix of high-quality genomic DNA extracted from six species: Sorghum bicolor (Great millet), Hordeum vulgare (Barley), Pleurodeles waltl (Iberian ribbed newt), Notamacropus rufogriseus (red-necked wallaby), Pteropus lylei (Lyle’s flying fox) and Nymphalis urticae (Small tortoiseshell).

To ensure optimal amplification of both primer pairs, another gradient PCR was performed using the same conditions as above but using the 6X_DNA sample as a template and the two primers mixed at a final concentration of 0.2 μM each. After several rounds of PCR optimization, we obtained two clear bands with the following settings: in a total volume of 20 μl using 1 μl of 6X_DNA (1 ng/μl), 10 μl of 2x QIAGEN Multiplex PCR Master Mix, 3.5 μl of primer mix (0.2 μM final concentration each primer), and 5.5 μl of ultrapure water. Thermal cycling conditions included an initial denaturation step at 95 °C for 15 min, followed by 35 cycles at 95 °C for 30 s, 55 °C for 90 s, and 72 °C for 90 s, and a final extension at 72 °C for 10 min. PCR products were visualized on an Agilent Bioanalyzer DNA 7500 Assay chip without further purification.

Finally, to test the sensitivity of our multiplex PCR, we used the aforementioned 6X_DNA control diluted in H2O and spiked into an eDNA sample in defined amounts ranging from 1 ng spiked to 0.001 ng in 10 ng eDNA (1/10000).

In order to increase the robustness of our results and extract the maximum amount of information from the DNA extracts, we opted to increase the eDNA input volume to 6.5 µl per replicate after confirming that this did not result in PCR inhibition (Ushio et al. 2022). Multiplex PCR amplification of eDNA extractions and the corresponding positive (1/10 dilution of 6X_DNA, 1 ng/μl) and negative controls (water and no-template controls, NTC) was performed in triplicate using the BR3 and 230_RFmod primers, as well as ITS2 UniPlant uniquely tagged to allow for individual sample and replicate identification. Final conditions for metabarcoding of plants and animals were as follows: 6.5 μl of extracted DNA, 10 μl of 2x QIAGEN Multiplex PCR Master Mix, 3.5 μl of primer mix (0.2 μM final concentration each primer) in a total volume of 20 μl. Thermal cycling conditions included an initial denaturation step at 95 °C for 15 min, followed by 35 cycles of 94 °C for 30 s, 55 °C for 60 s, and 72 °C for 90 s, and a final extension at 72 °C for 5min.

After PCR, amplicons were cleaned up and normalized in a single step following a bead-based normalization strategy (Suchan 2020; Hosomichi et al. 2014), reducing hands-on time and the need for tedious concentration-based normalization (Buchner et al. 2021).

SPRI beads (Sera-Mag, Cytiva) are diluted 1:20 in bead normalization buffer (2.5 M NaCl, 0.01 M Tris, 0.001 M EDTA, 0.05% Tween 20, 20% PEG 8000) to limit the binding capacity of the beads and recover a defined amount of DNA. To this end, the normalization solution was added to the sample at a 1:1 ratio (v/v), mixed with 1:1 (v/v) isopropanol, and incubated for 10 minutes at room temperature. Next a standard SPRI bead cleanup was performed, including two washes with 80% ethanol. For the final elution, the beads from twelve wells at a time were resuspended in a total of 50 µl PCR-grade water. These pools of twelve were subsequently combined into one pool and quantified using a Qubit fluorometer dsDNA High Sensitivity Assay Kit.

### Library preparation optimization and Oxford Nanopore Technologies (ONT) sequencing

A common problem when preparing a sequencing library of a pool of PCR products carrying unique identifiers is tag jumping or swapping (Esling et al. 2015; Rodríguez-Pérez et al. 2022). To minimize tag swaps during library preparation and false assignments of tags to samples during sequence demultiplexing, we adapted the Tagsteady protocol (Carøe and Bohmann 2020), initially established for Illumina sequencing, to Oxford Nanopore’s Ligation Sequencing protocol. To account for the higher sequencing error rate, we used longer 9 bp tags that can help discriminate different samples with greater confidence (Srivathsan et al. 2024).

Our modified library preparation protocol combines the Tagsteady End Repair step immediately followed by the Adapter Ligation and Clean-step from Nanopore’s Ligation Sequencing Amplicons V14 Kit (SQK-LSK114). By following this modified procedure, the tag-jumping effect is minimized while the total time for library preparation is reduced by skipping the clean-up step after End Repair, as in the Tagsteady protocol.

The final modified protocol to prepare ONT libraries using pools of individually tagged amplicons consisted of the following steps: End Repair was performed by adding 1.5 µl Klenow fragment exo-(5 U/μl), 2 µl T4 PNK, 0.5 µl ATP, 4 µl T4 DNA Ligase buffer (10X), 2 µl Reaction Booster, and 300 fmol amplicon pool. The reaction was incubated for 30 minutes at 37 °C followed by 5 minutes of heat inactivation at 65 °C. This product was directly used as input for the Adapter ligation and Clean-up step. From this point on, we proceeded as described in the ONT Ligation Sequencing Amplicons Kit (SQK-LSK114).

Each final library was run on an individual ONT Flow Cell (FLO-MIN114 R10.4.1) on a Mk1C or Mk1B device for at least 12 hours according to the manufacturer’s protocol.

### Long-range PCR

To set up a long-range metabarcoding protocol, we proceeded in a stepwise manner, from the high molecular weight DNA of a single species, to a mix of known species (6x DNA positive control) and finally to eDNA samples. At each step, we evaluated the size of the obtained amplicons and their concentration after PCR amplification using capillary gel electrophoresis (Bioanalyzer High Sensitivity DNA). Next, to transfer this to the field and integrate it with our already existing Multiplex PCR approach, we tested three different Polymerases: one Polymerase optimized for long-range PCR (PrimeSTAR, Takara), one engineered high-fidelity Polymerase (Kapa Hifi), and one highly robust Multiplex Polymerase (Qiagen Multiplex PCR).

The final protocol for amplification of long DNA barcodes for ribosomal DNA differed from the multiplex PCR protocol only in the thermal cycling conditions: initial denaturation at 95 °C for 15 min, 35 cycles of 94 °C for 30 s, 63 °C for 90 s, and 72 °C for 4 minutes, and a final extension step at 72 °C for 10 min.

### Barcoding Protocol

To perform insect barcoding experiments with our protocol, we set up a simplified version that skips the time- and cost-intensive step of DNA extraction. In brief, we extract DNA by boiling a part of a leg or wing in 10 µl of phosphate-buffered saline (PBS) for 5 minutes at 98 °C. We then subjected 2.5 µl of this crude extract to PCR amplification with either COI or rDNA primers following the protocols described above. After sequencing, these new barcodes can be added to the reference database. A similar approach can be employed for in situ plant barcoding by stabbing collected plant material with a pipet tip or a piece of fishing line (Lynagh et al. 2023). Samples from the Barcoding test experiment presented here used DNA previously extracted by the sample providers.

### Blanks and controls

Contamination leading to false detections is an important problem in eDNA-based biomonitoring, especially when using universal primers. While it is difficult to completely avoid contamination in a well-equipped molecular lab, it is impossible in a field-based scenario. Thus, the best practice is to use different types of controls like field blanks, extraction blanks, and finally PCR no-template controls (NTC). Where possible, DNA extraction and PCR amplification should be separated strictly into pre-PCR and post-PCR, with no flow of personnel or reagents from post-to pre-PCR. Before and after starting an eDNA experiment, all surfaces, pipettes, and machines should be decontaminated with appropriate reagents (de Silva Wijeyeratne and Gweon 2025). In addition, the use of several PCR replicates can increase confidence in detection when a taxon is identified across several replicates. Here we used all these approaches and advise doing the same when using our protocol.

A final step-by-step protocol including Multiplex PCR, bead normalization, Nanopore library preparation and sequencing is available (https://dx.doi.org/10.17504/protocols.io.eq2ly4dbplx9/v1).

### Computational methods

#### RTBioScan pipeline

We developed a bioinformatic pipeline that identifies taxa in real time with high accuracy. It is implemented in Nextflow DSL1 (Di Tommaso et al. 2017). Two run profiles are provided: the default environmental metabarcoding profile (RTBioScan^m^) and the voucher profile for single-specimen barcoding (RTBioScan^b^).

#### Metabarcoding pipeline

To enable real-time, high-accuracy species identification in environmental metabarcoding using Nanopore sequencing, the pipeline employs a three-layered basecalling strategy leveraging the computational trade-offs between Nanopore’s fast, high accuracy (hac), and super accuracy (sup) dorado models.

The analysis proceeds in rounds with a feeder process that monitors the MinKNOW output folder and repackages incoming POD5 files into chunks of 200,000 reads by default (each chunk is one round). Rounds are processed one at a time, and all the protocol (read pools, OTU definitions, consensus sequences, and reports) is carried forward per round, and results are updated cumulatively.

All reads in the available POD5 chunk of each round are initially basecalled with the fast model (without a quality filter) from the electrical signals recorded every 10 minutes (this can be specified in the MinKNOW configuration file). The resulting preliminary sequences are aligned with LAST against a screening database of 95% identity non-redundant of the used markers (COI and ITS2) from all kingdoms to discard off-target sequences and sequences targeting undesired taxa (Bacteria, Archaea, Fungi, etc.). A read is retained only if its first LAST hit is labeled for the right combination of marker and target kingdoms (COI–Metazoa or ITS2–Viridiplantae); all other reads (bacterial or fungal COI, fungal ITS2, rbcL, unaligned) are discarded. This step efficiently reduces dataset complexity and focuses resources on the target taxa. Taxonomic reports are updated during sequencing.

The recovered animal and plant reads (1-40% of all sequences, depending on sample type) are re-basecalled with the hac model, keeping reads with Q-score ≥ 10 (≥ 90% expected accuracy). Before marker screening, fast sequences undergo a global read length filter. Hac reads are trimmed and demultiplexed with cutadapt, requiring post-trimming marker-specific lengths (350 and 286 nt for COI and ITS2, respectively). Reads lacking a recognized tag combination can be retained without sample assignment only if they pass the primer-only rescue and length filters. Demultiplexed reads are clustered into Operational Taxonomic Units (OTUs) at 97% identity with CD-HIT (Fu et al. 2012), each OTU potentially representing one species. Although OTUs are rarely used with Nanopore data, clustering provides a natural ground for read selection and consensus generation (Srivathsan et al. 2024; Elbrecht et al. 2019).

Accordingly, we used an OTU classification to ensure the coherence of the assignments, reduce possible false positives, and refine the sequences from the most promising OTUs. Active OTUs are eligible for freezing after at least three recorded observations of the same representative sequence, when the sum of their recorded read counts is at least 50 and no successive count in the last three observations decreases by more than 25%. Frozen representatives are retained for incremental assignment at the configured OTU identity threshold (97% identity by default) with the CD-HIT incremental clustering protocol (Fu et al. 2012). Reads not assigned to frozen OTUs remain in an active pool that can be reclustered as new evidence arrives; clustering is reused when that pool is unchanged. Frozen representatives and consolidated consensuses retain selected results across rounds. This reduces repeated computation while preserving dependence on accumulated state.

Consensus consolidation uses a support rule within frozen OTUs (>= 10 reads and a representative Q-score >= 20). Consolidated consensus sequences may subsequently be reused from cache, skipping consensus recalculation. To control accumulated low-support evidence, RTBioScan applies configurable size- and assignment-based pruning policies, while preserving protected assigned evidence. The default environmental configuration used in this study includes removal of taxonomically unassigned reads and persistent (>=3 rounds) small (<= 3 reads) or unassigned OTUs.

Hac reads are compared to a reference database of barcode sequences using megablast (Zhang et al. 2000). Reference sequences were clustered at 98% identity, and representatives were labeled with the last common ancestor (LCA) of the taxa within each cluster. This limits assignments to the resolution retained in the representative’s lineage. Query assignments use the retained megablast target and identity thresholds of 98%, 95% and 92% for species, genus and family, respectively (Alberdi et al. 2018; Leray et al. 2013). Positive BLAST results are cached per read and reused across rounds.

For each OTU and sample, up to 50 new hac reads per round, ranked by their BLAST identity, are re-basecalled with the sup model, retaining reads with Q-score ≥ 15 (≥ 97% expected accuracy). Sup reads are cached and reintroduced in subsequent rounds, so that the number of sup reads per OTU accumulates over the run. A consensus sequence is generated per OTU and sample when at least 5 sup reads are available: reads are aligned with DECIPHER AlignSeqs and a consensus is called with ConsensusSequence. For consolidated OTUs, only reads with Q-score ≥ 20 are used. The consensus sequences of each sample are then clustered at 99% identity with VSEARCH to merge redundant OTUs and retrieve one representative per cluster, and the representatives are assigned taxonomically with megablast using the same database, parameters, and thresholds as above. Every round produces an updated HTML report and machine-readable tables (read counts per sample, replicate, and marker; OTU membership; consensus sequences and assignments). The end of the run is decided by the user from the saturation of detections in the report or from a pre-established sequencing schedule. Development of RTBioScan was assisted by Claude Code (Anthropic) and Codex (OpenAI); all code was reviewed and tested by the authors.

#### Detection criteria

RTBioScan applies read-level quality and length filters and OTU-level support filters, but it does not perform sample-level statistical filtering; it outputs tables of read counts and assignments per sample, PCR replicate, and marker. For the analyses reported here, we consider a taxon to be detected in a sample if 1) a consensus sequence was generated, 2) it was supported by more than five sup or hac reads, and 3) it was found in more than one PCR replicate. Taxa detected in any of the PCR-negative controls were excluded.

#### Barcoding (voucher) pipeline

Single-specimen barcoding (RTBioScan^b^) uses the same pipeline with the voucher profile, which adapts the parameters to samples dominated by one individual, in the spirit of recent large-scale Nanopore barcoding protocols (Srivathsan et al. 2024; Hebert et al. 2025). Read screening, hac basecalling, and demultiplexing are identical to the metabarcoding pipeline. OTUs are clustered at 99% identity; sup basecalling and consensus reads require Q-score ≥ 20, and consensus sequences are built from 10–100 sup reads counted over the whole cluster. Consensus sequences are clustered per sample at 99% identity as above, and the dominant cluster per marker and sample (highest read support) is exported as the voucher barcode, with the megablast assignment reported as a suggestion. For this process, a limited number of reads per sample are enough to obtain high-quality barcodes (Hebert et al. 2025).

#### Long-read rDNA pipeline

The rDNA metabarcoding pipeline is currently standalone and not part of RTBioScan. It is not a real-time analysis pipeline and is not implemented in nextflow, but rather an add-on at this point. Unlike RTBioScan, it starts from bascalled reads, in the best case dorado sup accuracy. The fastq files are demultiplexed by sample using cutadapt with the linked adapters option (Martin 2011). Reads are length- and quality-filtered according to specified parameters and clustered into OTUs using CD-HIT, as in the main RTBioScan pipeline. Next, consensus sequences are generated per OTU as in the main pipeline. This highest-quality sequence is used for rDNA annotation using ITSx as well as taxonomic classification using SINTAX search (Bengtsson-Palme et al. 2013; Edgar 2016; Rognes et al. 2016). By default, the Eukaryome v2 database is used for classification (Tedersoo et al. 2024). Classification was run for each of the identified rDNA regions (SSU/18S, LSU/28S, 5.8S, ITS1, and ITS2) individually and for the full sequence. A Consensus classification was generated as the lowest-level classification that agreed taxonomically with the other classifications. The pipeline is available at https://github.com/lewange/longread_rDNA_metabarcoding. Claude Code Sonnet 5 (Anthropic) was used to assist in developing the rDNA amplicon annotation pipeline and in analyzing the resulting data. All code was reviewed and tested by the authors.

### Construction of the COI and ITS2 database

The main public resource of barcode sequences is BOLD, which stores 23,883,262 sequences corresponding to 652,546 different species. Among them, the Cytochrome Oxidase I (COI) is the most used barcode for animals (19,973,552 sequences target its 5’ region for 442,223 species) and rbcL and the internal transcribed spacer 2 (ITS2) for plant species (195,292 and 149,023 sequences for 62,204 and 50,984 species, respectively). We have chosen our metabarcoding platform to target COI-5P and ITS2 because of their wide species coverage and taxonomic resolution.

Consequently, we started by recovering the animal COI-5P and plant ITS2 sequences in BOLD. Although BOLD is the most comprehensive resource of barcode sequences, the targeted regions have also been sequenced for many other species as part of non-barcoding efforts. To take advantage of this information, we recovered COI and ITS2 sequences from NCBI Nucleotide (https://www.ncbi.nlm.nih.gov/nucleotide/) with specific queries (*“(((((((cytochrome c oxidase subunit 1[Title] OR cytochrome c oxidase subunit I[Title] OR cytochrome oxidase subunit 1[Title] OR cytochrome oxidase subunit I[Title] OR COX1[Title] OR CO1[Title] OR COI[Title]))) NOT UNVERIFIED) “* for COI and “*((viridiplantae[Organism] AND its2) AND 100:10000000[Sequence Length]) NOT (uncultured OR environmental sample OR incertae sedis OR unverified)”* for ITS2), retrieving 3,787,982 and 250,544 sequences for 202,366 and 79,235 species, respectively.

COI records were integrated following the COInr/mkCOInr procedure (Meglécz 2023), which merges BOLD and NCBI records into a coherent taxonomy, restricts sequences to 100–2,000 nt with at most five consecutive internal Ns, excludes environmental, metagenomic, intron-containing, and non-COI records, and removes redundant substrings within taxa; non-metazoan records were excluded. ITS2 records were curated following Quaresma et al. (2024): removal of fungal sequences of non-ITS2 sequences (< 70% identity to a trusted vascular-plant ITS2 panel), of records with incomplete taxonomy and of Chlorophyta, and removal of intraspecific outliers for species with more than four sequences. Barcodes generated in this study (Table 1) were added to both collections.

**Table 1:** RTBioScan Barcoding results. Two plant samples and nine animal samples were used to test RTBioScan’s ability to generate high-quality barcodes. Assignment identity thresholds were species: 98%, genus: 95%, family: 92%. For unassigned specimens, closest reference matches were obtained with a separate exploratory search against the NCBI core nucleotide collection (core_nt) using BLASTN (standard, non-megablast parameters); these matches were not accepted as pipeline taxonomic assignments. Query coverage is reported separately from alignment length.

| Known species | Marker | Percent identity | consensus length | alined length | Taxon best hit | Rank assigned | Rank match |
| --- | --- | --- | --- | --- | --- | --- | --- |
| *Hordeum vulgare* | ITS2 | 99.7 | 292 | 292 | *Hordeum vulgare* | species | species |
| *Sorghum bicolor* | ITS2 | 99.7 | 292 | 292 | Sorghum | genus | genus |
| *Nymphalis urticae* | COI | 100.0 | 418 | 418 | Nymphalis | genus | genus |
| *Ursus thibetanus* | COI | 99.8 | 418 | 418 | *Ursus thibetanus* | species | species |
| *Pleurodeles waltl* | COI | 100.0 | 418 | 418 | *Pleurodeles waltl* | species | species |
| *Chamaeleo chamaeleon* | COI | 100.0 | 418 | 418 | *Chamaeleo chamaeleon* | species | species |
| *Pteropus lylei* | COI | 99.8 | 418 | 418 | Pteropodidae | family | family |
| *Pelecanus crispus* | COI | 99.8 | 418 | 418 | *Pelecanus crispus* | species | species |
| *Notamacropus rufogriseus* | COI | 98.3 | 488 | 460 | *Notamacropus rufogriseus* | species | species |
| *Varanus prasinus* | COI | 88.0 | 418 | 418 | Varanidae | unassigned | family |
| *Dracaena guianensis* | COI | 83.0 | 418 | 417 | Teiidae | unassigned | family |

Non-redundant reference database. To reduce the computational cost of sequence comparison and to make single-hit assignment robust to near-identical references, each collection was clustered at 98% nucleotide identity with CD-HIT, and each representative sequence was labeled with the LCA of the taxa in its cluster. Consequently, a species whose barcode is ≥ 99% identical to that of a congeneric species is represented at genus level (or higher) in the database, which prevents false species-level assignments at the cost of resolution for such species pairs. The resulting databases contained 898,771 (COI) and 70,544 (ITS2) representative sequences. Taxonomic lineages were retrieved with TaxonKit from the NCBI taxonomy (release 2024-06-24) and from BOLD-derived lineage tables for records without an NCBI taxid.

## Results

### Protocol setup and primer optimization

In order to enable detection of plants and animals in the same reaction, we first optimized PCR conditions for each of the primers individually and, after obtaining clear amplification for each of them, combined them into a multiplex PCR (Methods, Supplementary Figure 1). Next, we validated our Nanopore-based sequencing protocol against the published version of the Tagsteady protocol that uses Illumina sequencing. To this end, we compared four soil samples collected in Collserola Natural Park, Catalunya, processed with the original protocol and our version adapted to Nanopore sequencing. We detected both plant (ITS2) and animal (COI) taxa with similar sensitivity in both approaches (Supplementary Figure 2), supporting the technical compatibility of Nanopore sequencing with the Tagsteady protocol.

One big concern when using universal COI primers, as pointed out earlier, is a high fraction of bacterial reads. To alleviate this problem, we systematically optimized primer design in silico. We explored combinations of previously published primers (Elbrecht et al. 2019). For each combination, we exhaustively evaluated alternative primer sequences and new combinations of existing sequences for their specificity and sensitivity in animal detection based on their ability to recognize target regions in the set of complete bacterial and mitochondrial genomes available in the NCBI database and BOLD. In total, we tested 600 different primer pairs and selected the 10 most promising candidates based on this analysis.

To confirm the in silico predictions and select the best-performing primer pair, we tested them experimentally for their potential to specifically enrich for metazoan COI (Figure 2A). We see a clear improvement in 9 out of the 10 combinations over our previous primers (COI_1 (original), BF3/BR2), both regarding the number of detections and, importantly, the on-target reads. Based on these findings, we selected COI_3 consisting of the original reverse primer BR2 combined with 230_RFmod (Supplementary Table 1) for subsequent experiments. Next, we tested this new set of primers in comparison to the previously used BF3/BR2 primers multiplexed with the same UniPlant ITS2 primers on a range of different sample types (passive air and water filters, soil and insect tape). The high-content insect barcoding samples obtained more reads for both primer combinations and showed high on-target percentages for both COI and ITS2, likely explained by the high percentage of target DNA stemming from insects and pollen (Figure 2B). The biggest improvement can be seen for soil samples, where the COI on-target increased from ∼1% to more than 25% of reads. Smaller but decisive improvements can also be seen for water and air samples. Notably, ITS2 on-target percentages slightly increase as well in all conditions despite using the same primers, most notably in the insect tape samples (Figure 2C). This shows solid performance of both primer pairs in the multiplex PCR and a potential positive effect of improved COI primers for the ITS2 amplification that might be explained by primer interactions in the multiplex PCR reaction. Another possible explanation could be less competition for PCR reagents when a smaller amount of bacterial COI is amplified, which can also be seen as a relative increase of ITS2 target molecules over COI target molecules in the reaction.

**Figure 2.**
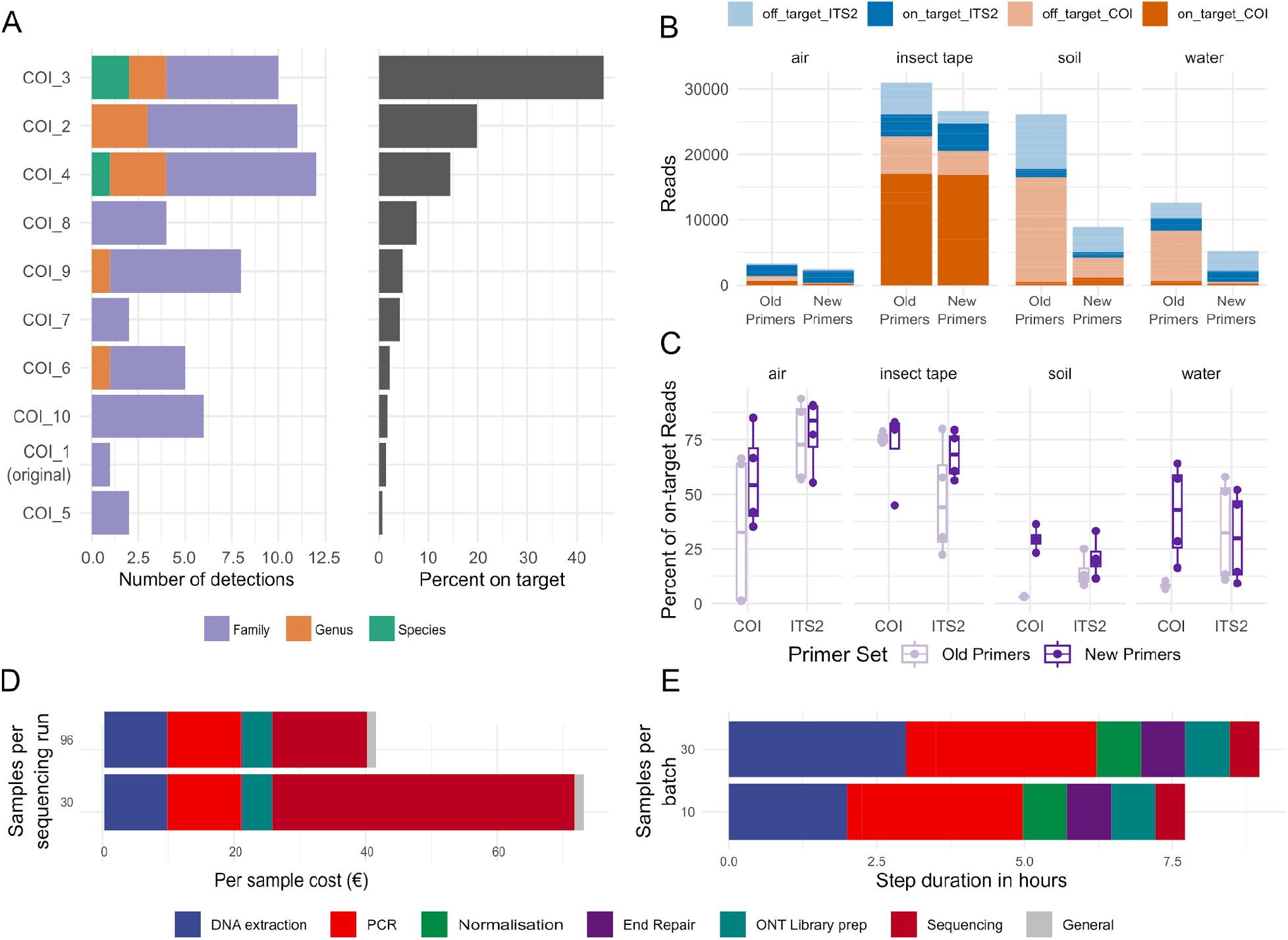
Experimental workflow establishment and validation. A) Experimental validation of primer specificity and sensitivity. Ten top candidate primer pairs were tested experimentally. The left barplot shows their ability to detect Metazoa at different taxonomic levels. The right plot shows the on-target percentages of each primer pair, i.e., the number of metazoan reads over all reads. B) Number of on-target and off-target reads for the original primer set and new primer set across sample types. C) Percentage of on-target reads for the old and the new primer set for ITS2 and COI, respectively, across sample types. D) Cost per sample including sequencing. Costs were calculated for 30 and 96 samples per sequencing flow cell, including 3 PCR replicates each. Colors indicate the contribution of each protocol step to the total cost. E) Processing time for 10 or 30 samples. Processing times represent the total processing time, including hands-on and hands-off times like incubations. Colors indicate the contribution of each protocol step to the total time.

We further tested the experimental limit of detection of our multiplexed enrichment PCR using a dilution series of known quantities of control DNA containing a mixture of 4 animal and 2 plant species. All PCRs showed successful amplification, and the presence of the expected positive control peaks was confirmed in all dilutions, including the 1/10000 dilution, corresponding to ∼ 0.17 picogram total DNA (assuming the six species contributed equal DNA mass, Supplementary Figure 3).

As one of the requirements for implementing eDNA biodiversity monitoring is efficiency, both in terms of cost and turnaround time, we calculated experiment timings and prices. Our final protocol can be performed for 30 samples in field conditions in just under 10 hours (Figure 2D) with a cost per sample including sequencing of 73 € (Figure 2E). While the PCR is one of the most time consuming steps it is mostly determined by the thermo cycling itself (hands off) and by cycler capacity in the field (in our case 3x 32 wells, 30 samples in triplicates plus controls). A well-equipped laboratory could scale up the number of samples easily without adding substantial processing time. The limiting factor for higher throughput is DNA extraction. With our current setup, we can process a maximum of 32 samples per extraction batch. Doubling the number of samples would double the time needed for DNA extraction. Being a bead-based extraction method, however, opens up avenues for processing more samples simultaneously in the future, and open science initiatives like bomb.bio make bead-based extractions accessible to scientists worldwide (Oberacker et al. 2019). The major cost factor in this protocol is the ONT sequencing flow cell followed by the PCR reagents and the DNA extraction. While we haven’t specifically tested the multiplexing capacity with our approach, sequencing more samples on the same flow cell is feasible and can drive down the cost per sample substantially (Hebert et al. 2025).

### Long-read metabarcoding

To harness the full power of Nanopore long-read sequencing, we implemented a long-range PCR approach for eDNA. To this end, we evaluated two sets of primers, one for the detection of animals covering a ∼8 kb region of the mitochondria (cytochrome B to COI) and one to detect plants and animals covering a ∼1-4 kb fragment of ribosomal DNA (18S to 28S) (Krehenwinkel et al. 2019).

While the rDNA fragment amplified robustly with all three tested polymerases, the mitochondrial fragment only amplified with the long-range polymerase and much less robustly. This can be attributed to differences in amplicon length and primer design. The mitochondrial primers are highly degenerate and slightly shorter than the rDNA primers, making them less well suited for long range PCR. In addition, the mitochondrial fragment is considerably longer, and inversions in the arrangement of the mitochondrial genes in some taxa complicate design. However, systematic optimization of primers is likely to yield successful amplification of a long mitochondrial barcode or the use of slightly shorter amplicons (Gajski et al. 2024).

It has been assumed historically that the majority of eDNA is highly fragmented; however, we found robust amplification across a range of different samples (soil, water, air, insect tape) using the ribosomal DNA primer spanning 18S to 28S of the rDNA, including ITS1, 5.8S, and ITS2 regions (Supplementary Figure 4). Analysis of rDNA fragments from soil samples showed broad amplification of Fungi, Viridiplantae, Metazoa, and other Eukaryotes. While Fungi and small plants dominated in this analysis due to the type of sample, this shows the general potential of these primers to work broadly across eukaryotes. (Supplementary Figure 5A). To ensure that the reads we obtained are intact rDNA reads, we annotated the sequence features using ITSx, confirming that the majority of sequences were intact (Supplementary Figure 5B and C). In line with this, we compared the sequence lengths across domains and observed consistent length variation as expected from this region (Supplementary Figure 5D).

### RTBioScan: a pipeline for real-time biodiversity analysis

The development of our real-time analysis pipeline was driven by a need to optimize computing performance while at the same time getting high-quality taxonomic assignments. While the fast model of the Nanopore dorado basecaller can keep up with the speed of sequencing, the more accurate high accuracy (hac) and super accuracy (sup) models are too computationally intensive and thus slow down processing markedly (Figure 4 A and B). To overcome this bottleneck, we developed an approach that dynamically selects the most informative reads for high- and highest-accuracy basecalling. This approach is illustrated in Figure 3 and roughly follows a taxonomic hierarchy. In the first stage, all reads are basecalled with the fast model and assigned to a marker and kingdom using LAST; off-target reads are discarded. The remaining reads are basecalled with the hac model, demultiplexed by sample and marker, clustered into OTUs, and pre-assigned taxonomically by megablast. Then, up to 50 reads per OTU and sample in each round are basecalled with the sup model. Finally, 5-15 sup reads per OTU are aligned and a consensus barcode (Hebert et al. 2025) is generated, which is reassigned taxonomically.

**Figure 3.**
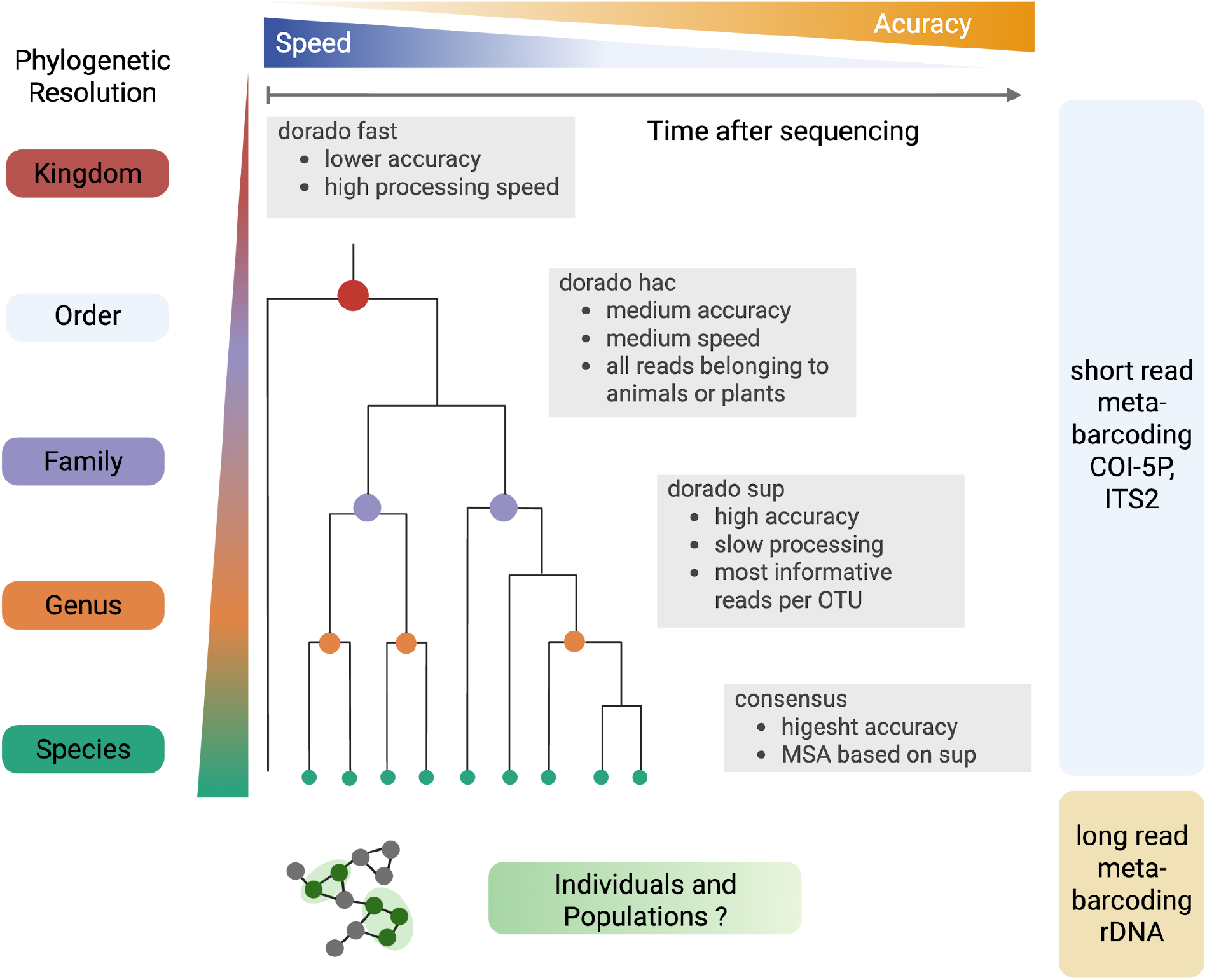
RTBioScan read sampling strategy. Schematic overview of the hierarchical dynamic read sampling employed by RTBioScan. RTBioScan selects reads for high and highest-accuracy basecalling based on their informativeness. This loosely follows a taxonomic hierarchy, reflecting the increasing resolution in taxonomic assignments of reads of higher quality. First, all reads are basecalled in the fast model of dorado to filter out reads not matching either plants or animals. All reads passing this filter are basecalled in dorado’s high accuracy mode (hac). In subsequent steps, reads are clustered into OTUs and selected based on their value for highly accurate taxonomic assignments to be basecalled with the super accuracy (sup) model. Selected accumulated high-quality reads are used to generate or reuse an OTU consensus, which is then compared with the reference database; frozen OTUs and consolidated consensus sequences are carried forward so that later rounds process only new reads.

**Figure 4.**
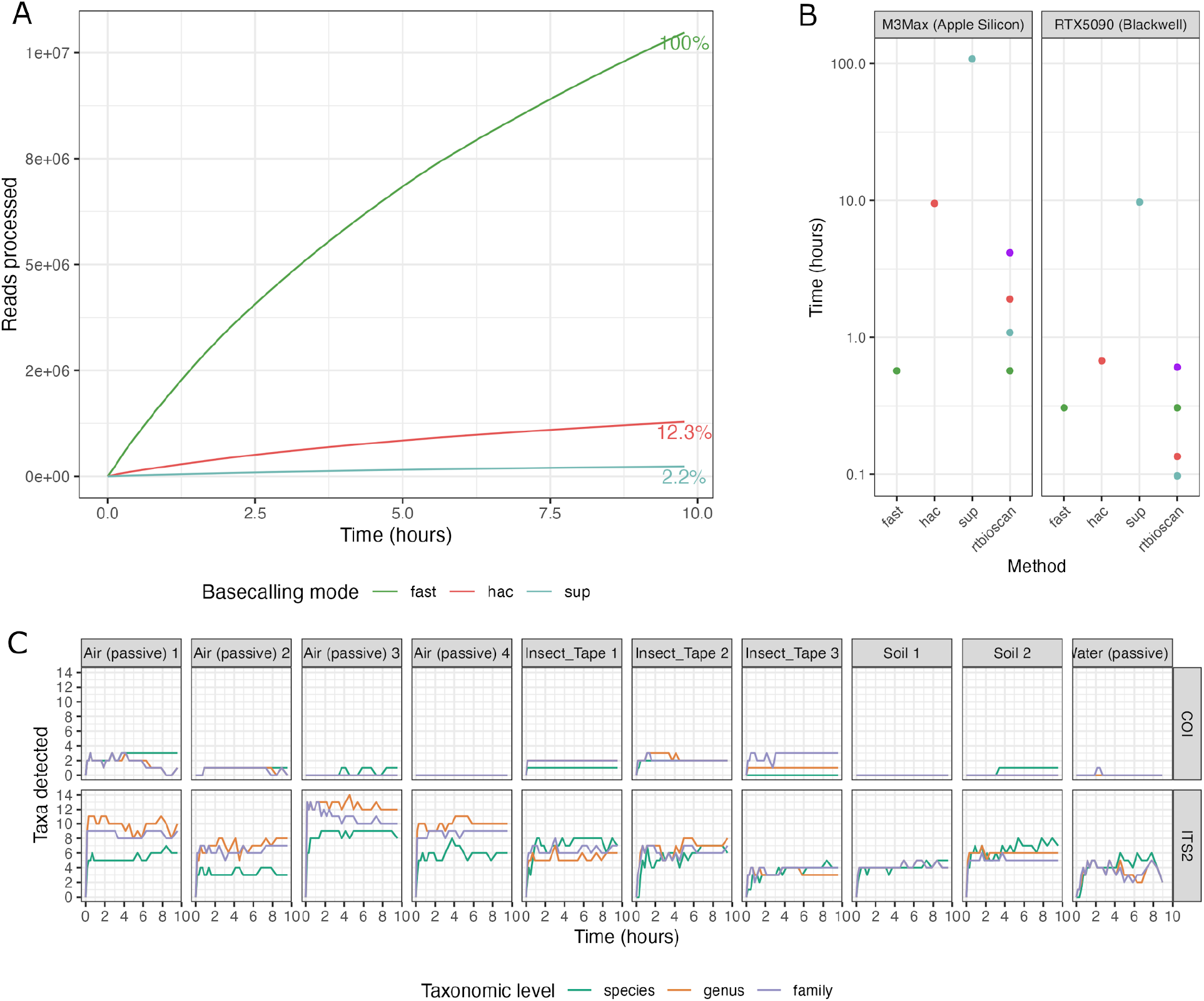
RTBioScan is computationally efficient and can monitor taxonomic identifications in real time. Reads basecalled by RTBioScan and taxonomic detections over time. A) All reads are basecalled in fast mode, while only 12.3% were basecalled in hac and 2.2% in sup mode. B) Basecalling times for 10 million reads with the three different models and with the RTBioScan read sampling strategy based on the read fractions in panel A. Performance was tested on two state-of-the-art GPUs, Apple Silicon’s M3Max and Nvidia’s RTX5090. C) Taxonomic detections at the species, genus, and family level over the time of sequencing for 10 different samples and four sample types.

Using this strategy, the RTBioScan pipeline allows taxonomic assignment in real time, identifying species based only on highly accurate sequences. Despite the overhead of basecalling individual reads up to three times with the different models, RTBioScan produces sup quality reads in a time range close to or faster than running the hac model on the entire set of sequences (Figure 4B). This shows that high-quality real-time basecalling is feasible already when using dynamic read sampling strategies as employed here.

With this real-time analysis, we can monitor the increase in detections at different taxonomic levels over the course of sequencing. While the samples sequenced here differ drastically in complexity, we observed a very fast saturation after 2-3 hours of sequencing in all samples, suggesting that additional multiplexing capacity may be available (Figure 4C). From a performance optimization point of view, our real-time survey of species identification allows us to determine the saturation point for our sequencing runs, optimizing the usage of the sequencer flow cells and avoiding the waste of time and sequencing resources in low-diversity or contaminated samples to focus the sequencing in more informative samples. The RTBioScan pipeline automatically produces a report that functions as an automatically updating sequencing dashboard (Supplementary File 1). With rounds of 200,000 reads and the hardware used here, RTBioScan updates its report, providing an estimation of the species present every ∼30 minutes; the interval scales with sequencing throughput and the configured round size.

Importantly, this optimization of sequencer usage allows increasing the number of runs per sequencer per time and per budget.

### An example of real-time biodiversity monitoring

To showcase the power of our approach, we processed 25 samples in triplicate belonging to four different sample types using our unified protocol. The number of reported taxonomic detections tended to stabilize during the first 2–3 hours under the tested assay and analysis settings. (Figure 4 C). Although this does not imply that all taxa in the samples were recovered, it provides a practical indicator of a drastic reduction in additional taxonomic detections. This saturation analysis is useful to decide when to stop a run (unless we aim to detect species with low DNA levels in very long runs). We saw the highest number of plant identifications in the air samples, and most animal identifications in the insect tape samples that physically trap insects (Figure 5A). Across all samples, we detected a total of 47 plant species (genus: 46, family: 51) and 11 animal species (genus: 11, family: 13). ITS2 detections cover a wide range of plant taxa known to be present in the sampling area, like pines, oaks, and oil trees (Figure 5B and C). Animal detections were less frequent than plant detections in these samples. This could reflect low template availability from sampling, amplification, computational filtering, and reference-coverage effects. Good detection of the positive controls shows the general ability of our primers to detect animals, and our sensitivity test shows that they are also efficient in detecting minute quantities of metazoan DNA, suggesting it could be due to a true absence of metazoan DNA in these samples (Supplementary Figure 3).

**Figure 5.**
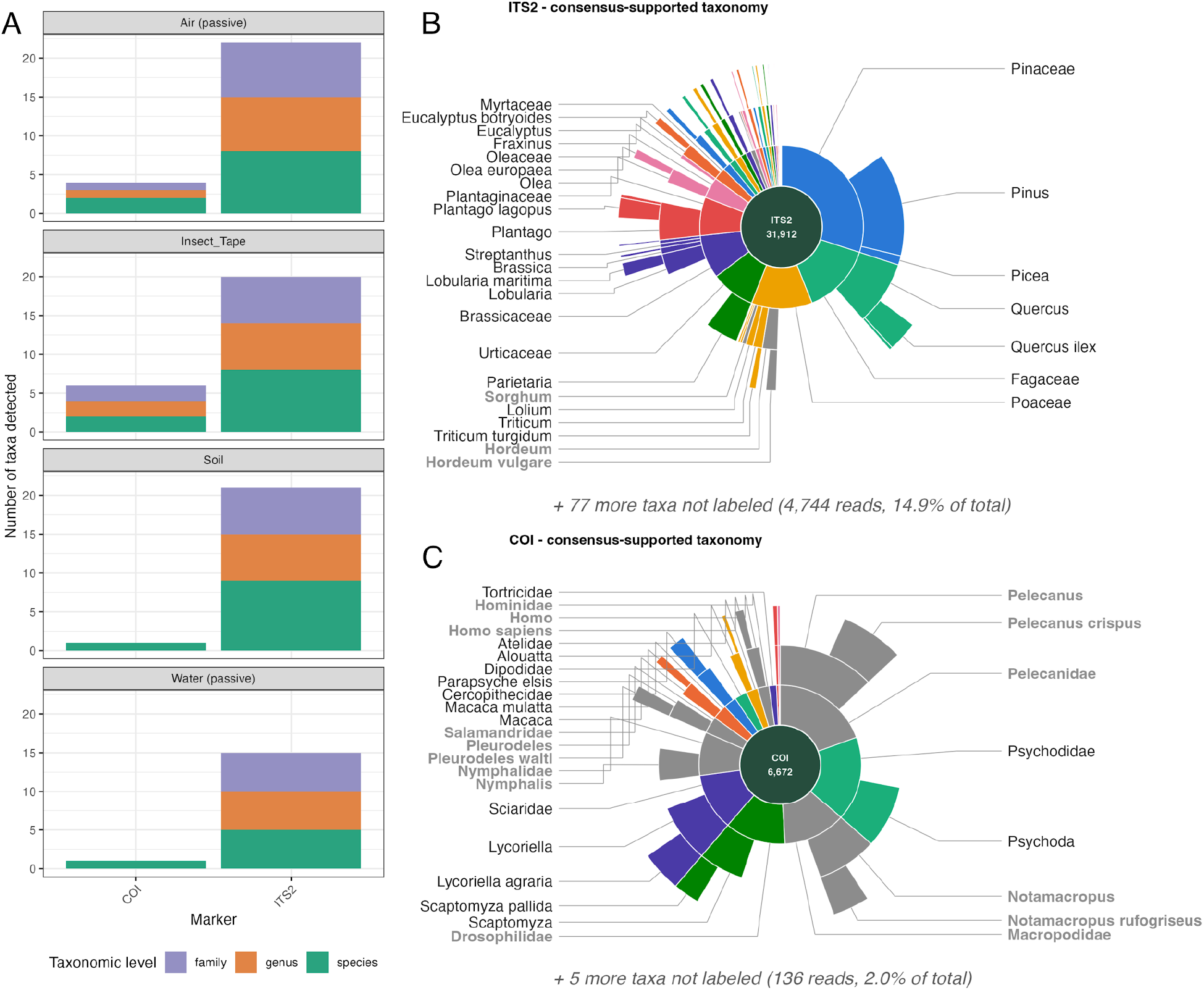
RTBioScan results for universal plant and animal metabarcoding across sampling methods. A) Number of taxonomic assignments at different levels (species, genus, family) and across sampling methods for COI and ITS2. B) Sunburst plot of ITS2-based identifications across all samples. C) Sunburst plot of COI-based identifications across all samples. Greyed-out areas correspond to positive control samples or contaminations. The area of the wedges corresponds to the number of reads.

We also detect low levels of contaminant primate DNA (Atelidae, Alouatta, Cercopithecidae and Macaca) likely stemming from processing samples in the same laboratory used for extractions of primate DNA. These are particularly found in the passive air samples, likely due to the low amount of DNA extracted from this type of sample. We see this despite taking all the typical precautions and using controls as outlined in the methods section Blanks and Controls. This highlights the risk of universal primers like the ones employed in this study, as they are susceptible to even the lowest levels of contamination and urges us to take the strictest precautions, especially using field blanks, extraction blanks, and PCR negatives.

### Integrated single specimen Barcoding

While metabarcoding is a powerful tool for biomonitoring, a main challenge, especially in undersampled remote areas and biodiversity hotspots, is the incompleteness of barcode databases (Luypaert et al. 2026). To enable researchers to perform barcoding and metabarcoding using the same reagents and protocol, we validated our protocol’s performance for single specimen barcoding. In addition, we added functionality to the RTBioScan pipeline to run in voucher mode (RTBioScan^b^) (see Methods). As a proof of concept, we sequenced nine animal and two plant samples of known identity. Consensus barcodes yielded six species-level, two genus-level, and one family-level assignments concordant with the expected specimen taxonomy; two specimens remained below the assignment threshold. Species-level hits had 98.3–100% identity. These comparisons assess concordance with reference records and do not independently estimate consensus sequence error.

Despite our threshold for species assignment being 98%, some samples matching entries in the database almost perfectly >99.7% were only assigned at lower taxonomic levels. This is expected from the database construction: barcodes clustered at 99% identity are represented by a single sequence labeled to their last common ancestor, so species with near-identical barcodes (e.g., Sorghum spp.) are assigned at genus level even when the query matches a reference at 99.7%. Two particularly interesting cases are the two reptiles, *Varanus prasinus* and *Dracaena guaiensis*. Their closest BLAST hits match at the genus/family level with the expected taxonomy but are not assigned to a specific rank due to the identity thresholds employed. This could potentially highlight more variability in the barcoded region for reptiles and raises issues when using strict identity thresholds. In summary, we show that Nanopore-based barcode generation alongside metabarcoding can be an additional tool in the field, making this pipeline an important addition to close gaps in existing reference databases.

## Discussion

Building on the highly efficient and reliable Tagsteady protocol and the NanoRTax pipeline, we aimed to design a comprehensive sample-to-species identification workflow. We systematically optimized both the experimental and computational protocols to reach higher specificity, sensitivity, and efficiency. This included rigorous testing of various PCR protocols, primer combinations, and computational pipelines.

As a result of the computational and experimental evaluation of 600 primer combinations of known primers and modifications thereof, we combined the reverse primer BR2 with a modified version of 230_R (Elbrecht and Leese 2017). Although 230_R was originally designed as a reverse primer, we use its reverse complement as a forward primer here and introduce additional degeneracy to enhance sensitivity. This combination achieved approximately 40% *in silico* and experimental specificity for animal identification (∼60% bacteria) compared to the 1% (99% bacteria) obtained by BF3 and BR2 (Figure 2 B, C). However, this specificity might come at the cost of species detection sensitivity when evaluated using samples with low bacterial contribution (mock community-like samples such as insect traps). As a result, we suggest this pair of primers (BR2 and 230_RFmod) to address animal detections in complex environmental samples like water, soil, or air, while favoring the original pair of primers for barcoding purposes or samples highly enriched in animal DNA.

To detect plant and animal species at the same time, we used a Multiplex PCR, combining ITS2 and COI primers in the same reaction. Highly multiplexed PCRs have been used for population genetics and individual identification but have not been extensively explored for metabarcoding thus far (Campbell et al. 2015; Hayward et al. 2022). In the future, we aim to scale this approach to more markers, for example, including 12S and 16S for animals and matK and rbcL for plants. It has been shown that the combination of multiple markers can help to overcome gaps in species representation in Barcoding databases (Luypaert et al. 2026). One concern with this approach is that some primer pairs will work more efficiently than others in the mixture, reducing the amount of information we can obtain compared to the singleplex. While we see relatively even representation of COI and ITS2 reads in our libraries, this should be explored further in the future, especially when combining more markers. On the other hand, this makes more efficient use of low-concentration samples by being able to increase the amount of DNA per reaction.

Our final protocol combines bead-based eDNA extraction, streamlined library preparation with SPRI-bead normalization, and fast ligation-based library preparation. Requiring only minimal equipment, i.e., a portable thermal cycler, a microcentrifuge, and magnet racks, this protocol enables the metabarcoding of 30 samples at a cost of 73€ per sample. It delivers a preliminary biodiversity assessment within 10 h (9 hours processing and 1 hour sequencing).

Further optimizations have the potential to make this protocol even more efficient, robust, and portable. In particular, using an isothermal reaction instead of PCR avoids amplification bias and doesn’t require a thermocycler (Plewnia et al. 2025). In addition, click chemistry-based Nanopore library preparation using dibenzocyclooctyne-carrying primers can further reduce library preparation time and avoid ligation-derived chimeric reads (Łypaczewski and Shapiro 2026). Recently, shotgun sequencing combined with genome skimming has been used to enable biodiversity monitoring without pre-selecting certain regions or taxa (Nousias et al. 2025; de Medeiros et al. 2025; Sullivan et al. 2025). While this is a very powerful approach, it is extremely costly and still relies on species representation in genome databases. With global initiatives to generate reference genomes for all vertebrates, animals, and more generally all extant species underway, this will become less of a problem in the future (Formenti et al. 2026; Lewin et al. 2018). However, currently making use of the vast existing barcoding databases is the better strategy, especially in less intensively studied regions.

Longer barcodes can provide more information about intra-species variability (Gajski et al. 2024). Here we use a ribosomal DNA barcode for metabarcoding and show that long fragments (3.5-4 kb) can be obtained from environmental samples. This could enable us in the future to go further than species-level detection and to distinguish different populations or even individuals. Recent studies using short DNA barcodes have shown that they can be used to reconstruct haplotypes and link intra-species variation to sampling location (Riley et al. 2025).

This application of DNA barcodes is also called metaphylogeography and will enable the field to move from species inventories to spatial abundance measurements (Serrana and Watanabe 2023; Turon et al. 2020). With longer sequences, the resolution and informativeness of these approaches increase. However, there are two important caveats for the proper adoption of long-range PCR in the field. In particular, for nuclear rDNA, intragenomic copy variation, paralogy, and pseudogenes can complicate interpretation. Mitochondrial markers face a different confounder, amplification of nuclear mitochondrial insertions (NUMTs). (Gajski et al. 2024). Second, similar to metabarcoding, better and more accurate databases will be needed in order to classify species and individuals with all the complexities of haploidy in nature. We are proposing a molecular protocol here to establish this methodology as a proof of principle, but an in-depth exploration of better markers and better bioinformatic solutions will still be needed to properly deploy this technology in the field for individual recognition. In general, by overlapping long barcodes with common barcoding regions like COI, cytB, and ITS2, it is possible to make use of existing databases for taxonomic identification and subsequently extract additional information on interspecies variability from the whole sequence.

One of the most significant advancements of our workflow is the real-time processing capability of RTBioScan. Leveraging the portability and high-throughput nature of Oxford Nanopore Technologies (ONT) MinION sequencers, RTBioScan processed over 1 million reads per hour while sequencing continued, delivering updated species identifications based on sup quality consensus sequences of the reads being generated within one round (∼30 min on a MacBook Pro with a M3Max GPU). This allows for near-instantaneous species identification with super/high accuracy basecalling, in the field and without the need for internet access or high-performance computing facilities. As for the molecular protocol, this aspect is highly relevant in regions where laboratories don’t have expensive HPC clusters at their disposal.

Like the molecular protocol, the RTBioScan pipeline can be used for metabarcoding as well as for single specimen barcoding using the “voucher” mode. This feature allows switching between these two applications seamlessly with the same exact protocol, reagents, and equipment. In the future, we plan to combine the detection of new barcodes and the analysis of complex mixtures with metabarcoding by automating the supplementation of the database with the newly generated barcodes in real time, which implies re-running the non-redundant clustering and LCA labeling. The nature of our broad phylogenetic targeting leads to a high number of OTUs that can currently not be identified at the genus or species level due to database gaps. However, the fast, reliable nature of our pipeline makes periodic reanalysis of existing data with an updated database feasible to identify these unknown taxa.

Our approach is designed for the broad application of eDNA to monitor biodiversity across a large area and continuous sampling over long periods of time. By designing a complementary real-time, high-accuracy analysis pipeline, we provide an integrated workflow for biodiversity monitoring in remote locations. These developments ensure that our protocol is scalable for broader application in real-world settings.

In conclusion, our universal plant and animal eDNA detection workflow provides an efficient approach for biodiversity monitoring. By addressing the limitations of traditional eDNA methods and integrating current methodological developments, we offer a versatile, real-time, and field-applicable tool for biodiversity assessment. This solution demonstrates potential for application in ecological research, conservation, and environmental management.

## Acknowledgements

We are grateful to the Laboratory of Bioacoustics (LAB) at the Universitat Politècnica de Catalunya, led by Michel André, for giving us the possibility to participate in the Xprize Rainforest competition, which inspired the development of this workflow. We thank the Barcelona CRYOZOO as well as Roger Vila for providing samples for the assay establishment. This work was funded by Sabine Plattner African Charities (SPAC).

---

**Supplementary Figure 1.**
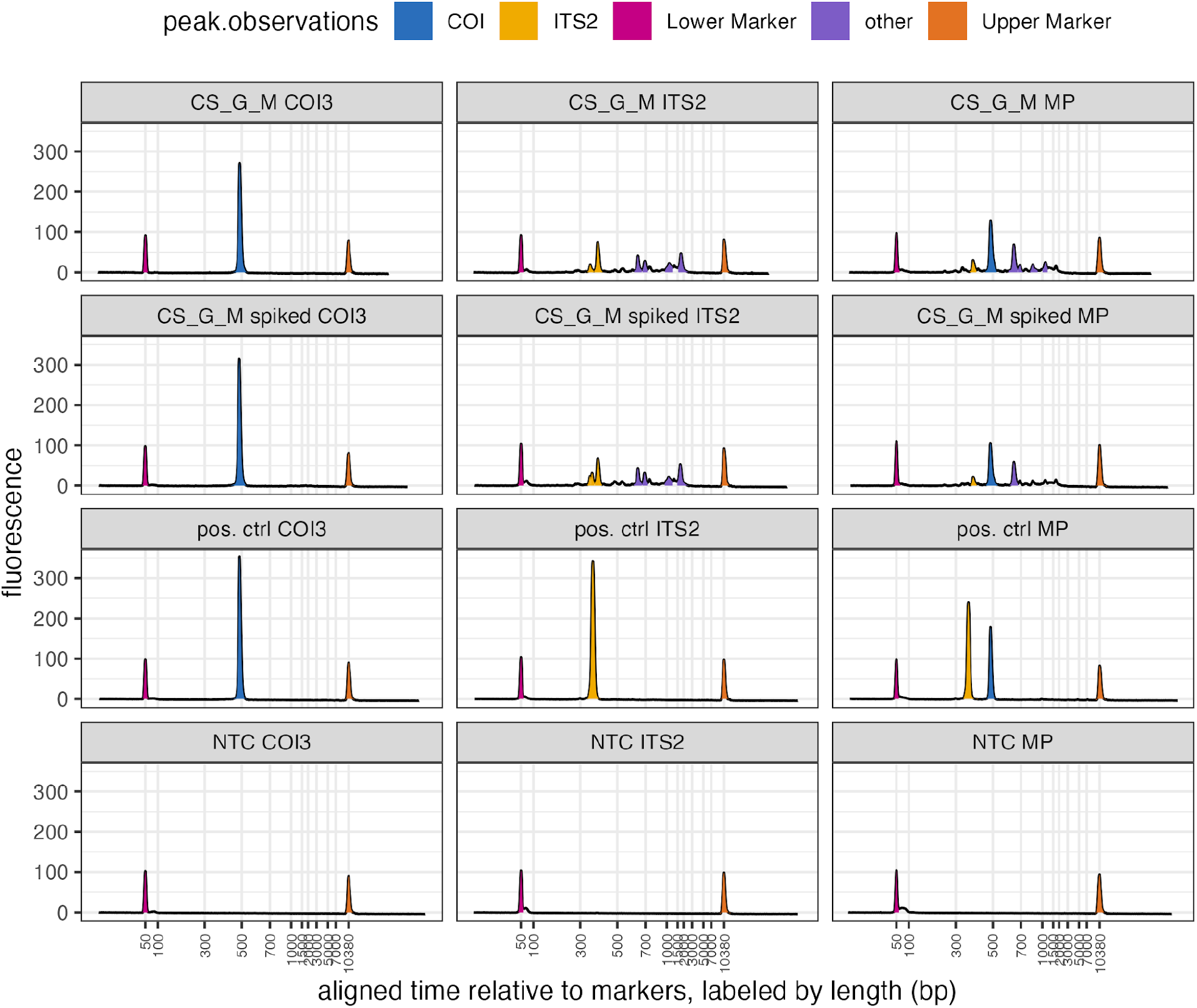
Comparison of Singleplex and Multiplex PCR. Bioanalyzer profiles of the individual primer pairs (COI, ITS2) and the multiplex PCR. Two eDNA samples from soil (CS_G, CS_G_spiked), as well as the positive control 6X_DNA, all show discrete patterns for the individual primers and a combination of the profiles when amplified in one reaction using multiplex PCR. The no-template controls (NTC) show no significant formation of primer dimers when both primers are combined.

**Supplementary Figure 2.**
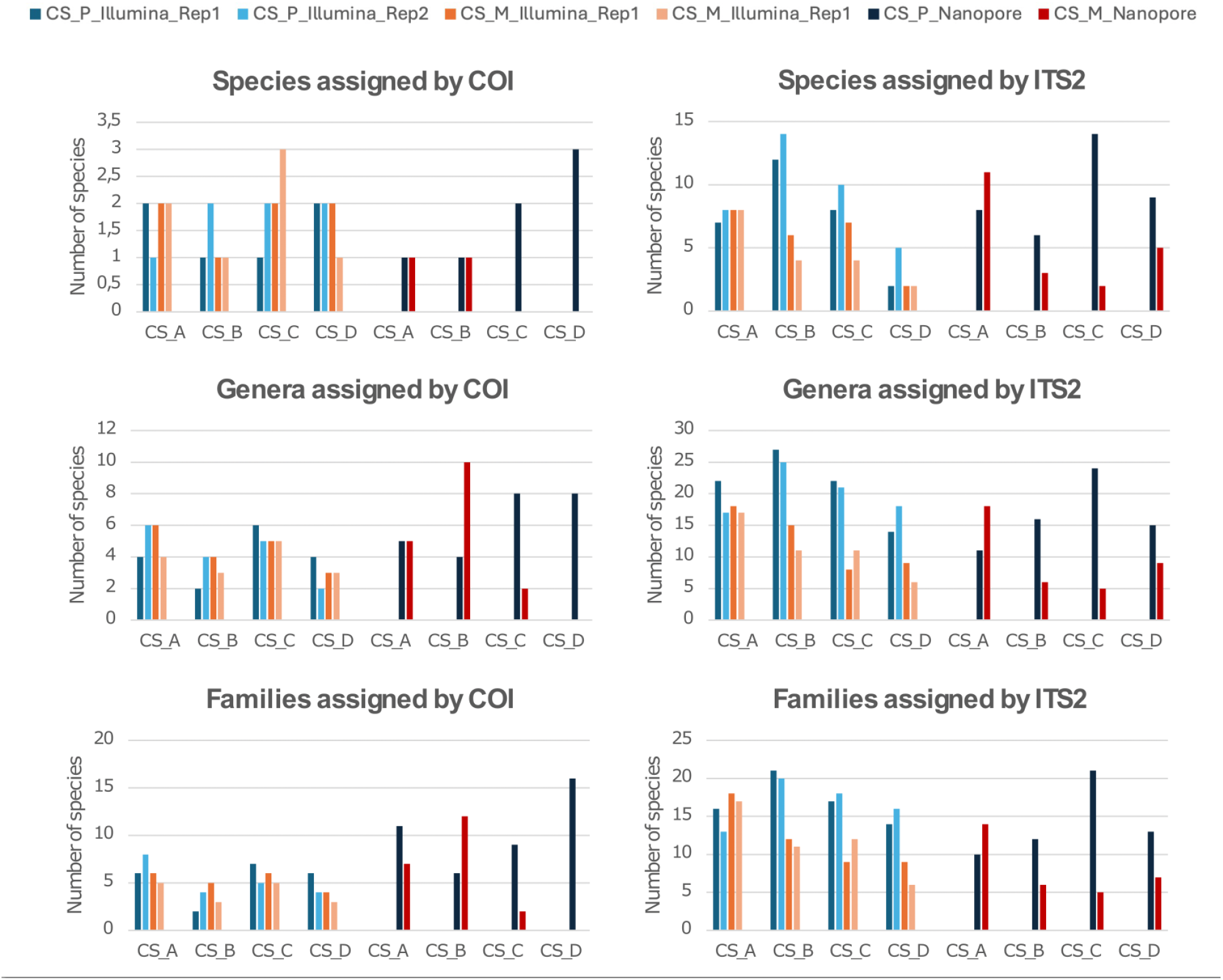
Tagsteady using Illumina and Nanopore sequencing. DNA of four soil samples (CS_A, CS_B, CS_C, CS_D) was extracted using the Qiagen PowerSoil (P) or Zymo MagBead (M) kit and sequenced on an Illumina MiSeq instrument or an ONT MinION (R10.4).

**Supplementary Figure 3.**
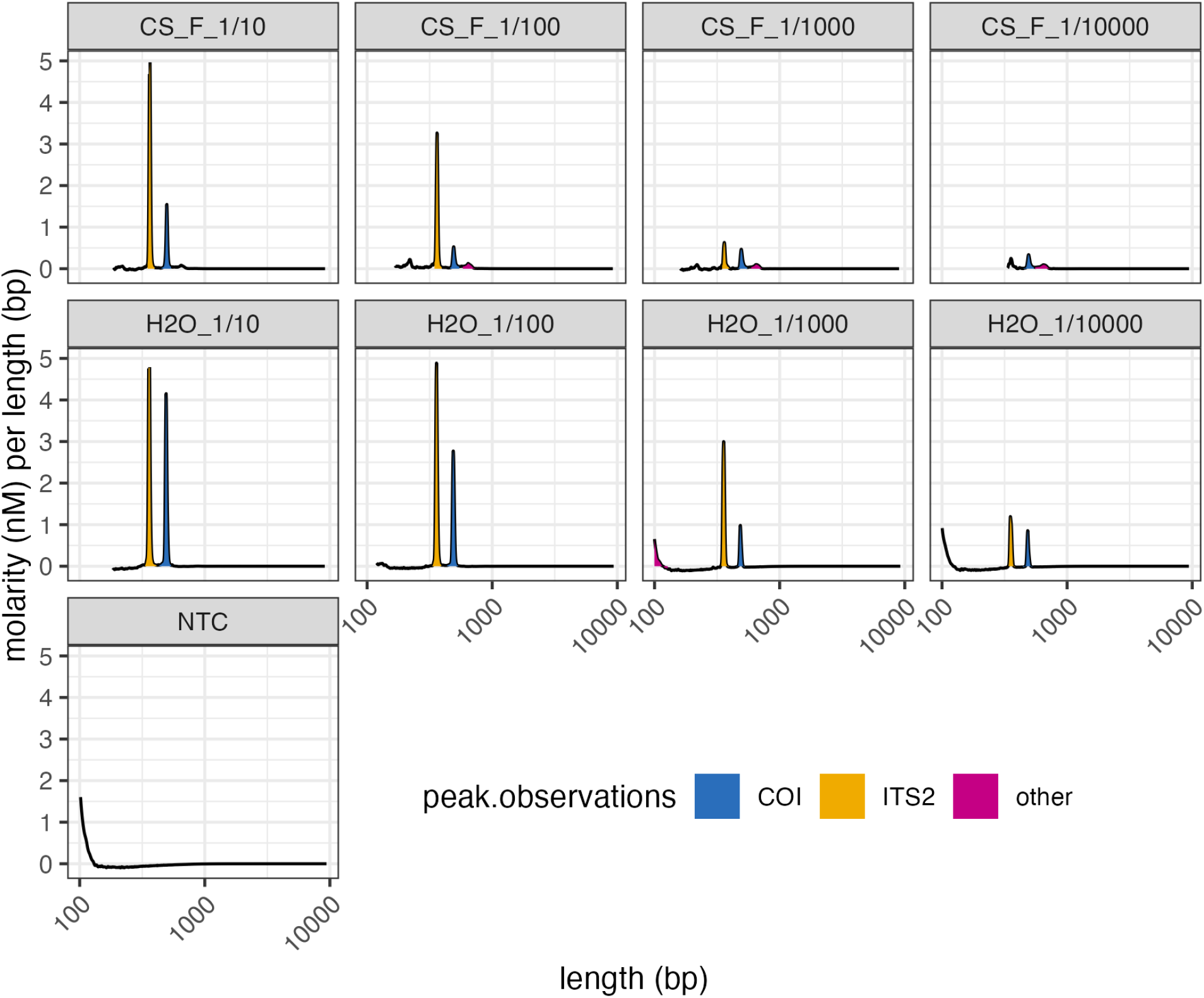
Sensitivity of the Multiplex PCR. Bioanalyzer profiles of defined input amounts to test sensitivity of amplification. The 6X_DNA control was diluted in H2O and spiked into an eDNA sample (CF_F) in defined amounts ranging from 1 ng to 0.001 ng spiked into 10 ng eDNA (1/10000). The control alone, diluted in H2O, was also tested separately and resulted in amplification even at the lowest concentration.

**Supplementary Figure 4.**
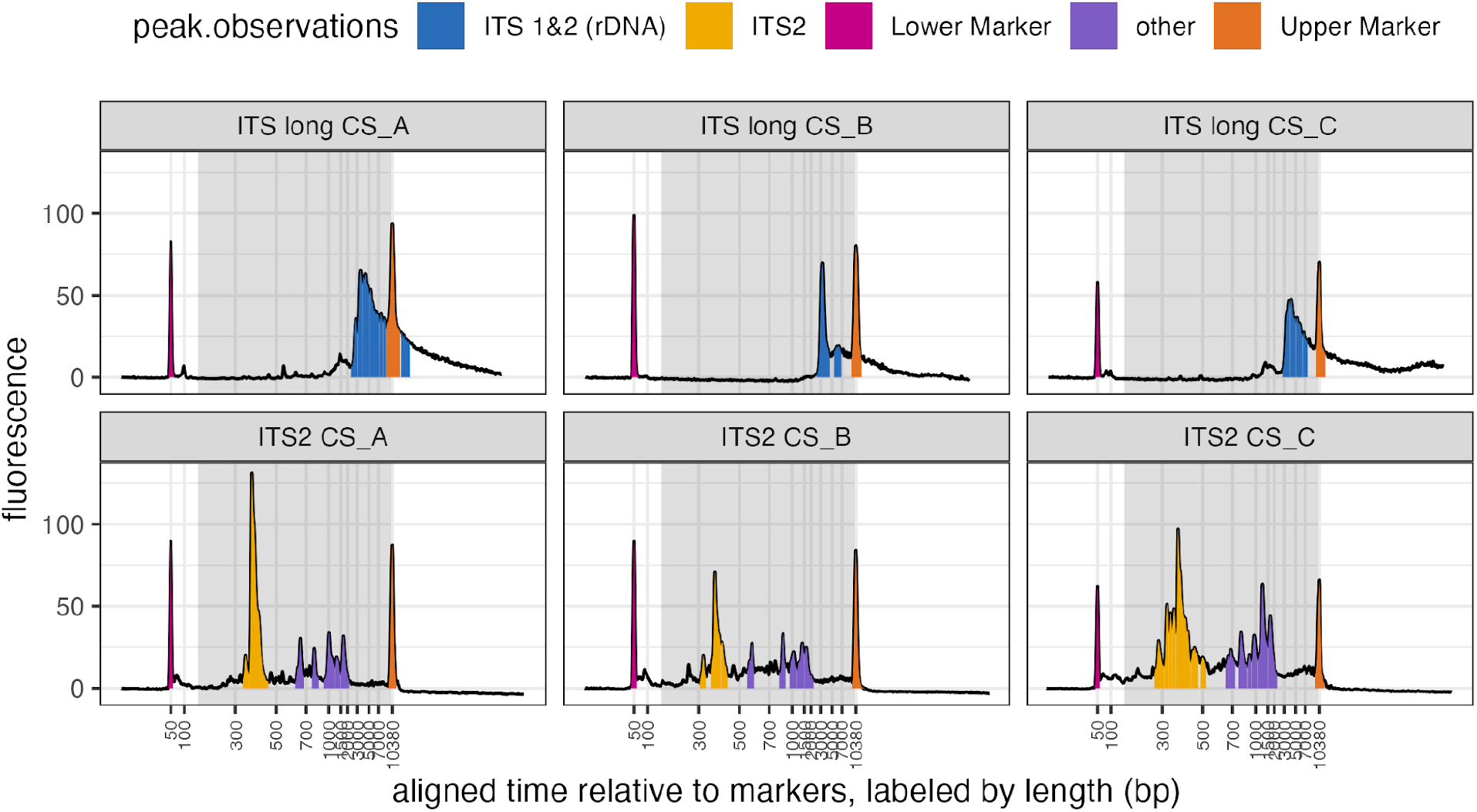
Comparison of amplification of a short ITS2 amplicon and a long rDNA amplicon. Bioanalyzer profiles of three samples CS_A, CS_B and CS_C with ITS primers and with rDNA primers. Expected amplicon size ITS2 187–387 bp (Moorhouse-Gann et al. 2018), rDNA/ITS long 3200 - 4500 bp (Krehenwinkel et al. 2019)

**Supplementary Figure 5.**
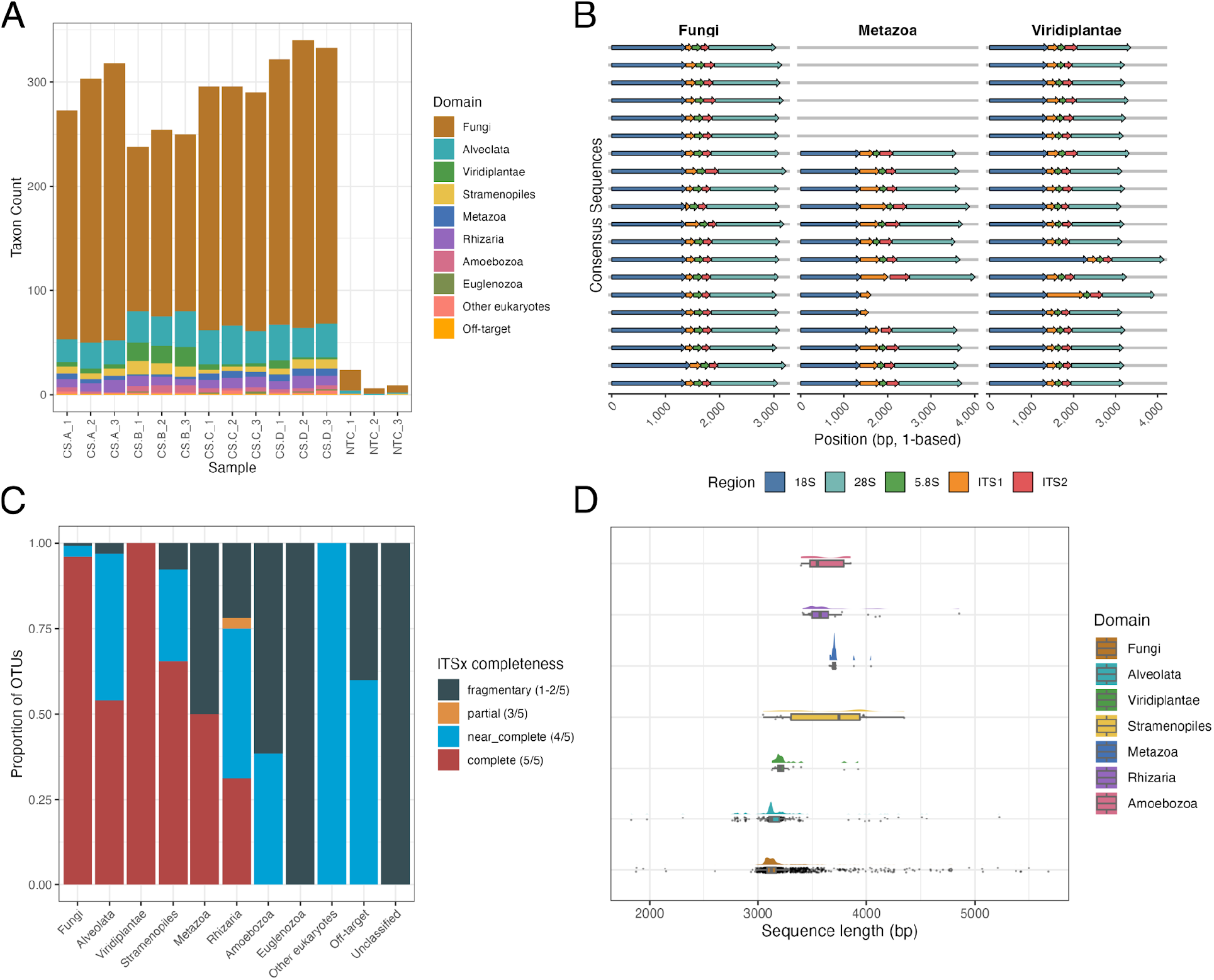
Long-range PCR enables rDNA-based metabarcoding. A) Taxonomic assignment using long rDNA amplicons. B) Structure of representative rDNA reads. C) Intactness of recovered rDNA by domain. D) Read length of sequenced amplicons by domain.

## Supplementary Tables

**Supplementary Table 1:**
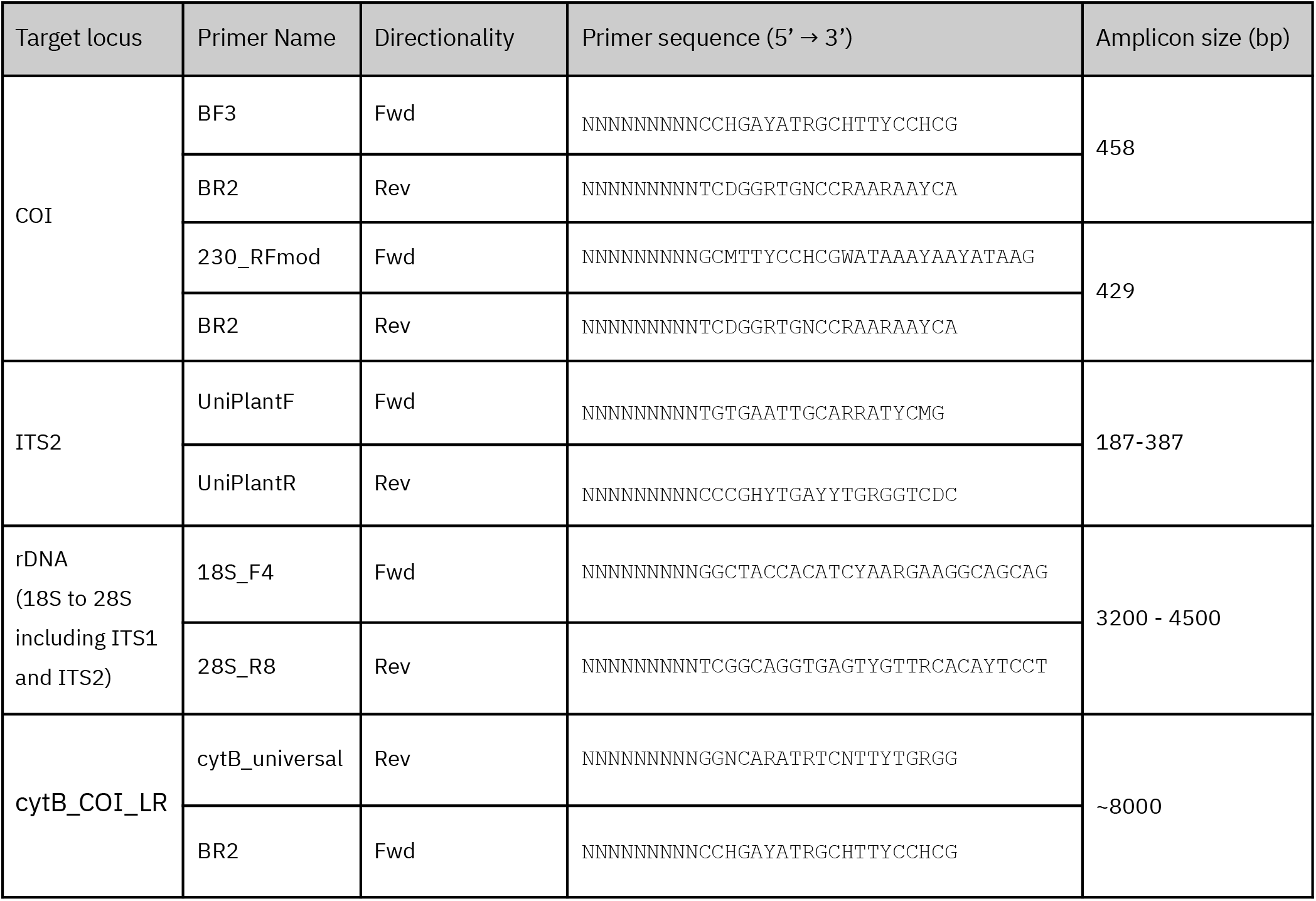
Primers used for metabarcoding. A 9-nucleotide-long tag is added to the 5’ ends of forward and reverse primers to allow for individual sample identification after sequencing.

## Data Accessibility and Benefit-Sharing

Both the experimental as well as the computational protocols and the barcoding database are publically available. A step-by-step protocol is available on protocols.io (https://dx.doi.org/10.17504/protocols.io.eq2ly4dbplx9/v1). The RTBioScan pipeline is available on GitHub: https://github.com/funcomicscnb-cpu/RTBioScan. The rDNA analysis pipeline is available at https://github.com/lewange/longread_rDNA_metabarcoding. Raw Nanopore sequencing data has been deposited at ENA.

## Author Contributions

ELG, DJ and TMB devised the experimental and computational strategies, developed the initial protocols and supervised the project. LEW, JPM and PAM tested and optimized the laboratory and computational methods. DJ implemented the main pipeline and the metabarcoding mode. JPM implemented the voucher mode and export. LEW implemented the rDNA long-read analysis, prepared the figures, and wrote the manuscript with input from all co-authors.

