## Supplementary Data 1 for "A sample-to-identification workflow to monitor animal and plant biodiversity using metabarcoding and real-time analysis": report.html

RTBioScan Run Info (run TS\_May2)


Back to index

### RTBioScan Run Info (run TS\_May2)

Last update: **May 19, 2026 11:24:14 UTC**

#### Global Overview

Sample Details
Rounds Info
Additional Info

#### Sample Details

Global Overview
Rounds Info
Additional Info

#### Rounds Info

Global Overview
Sample Details
Additional Info

#### Additional Info

Global Overview
Sample Details
Rounds Info

#### Warnings

×

##### Figure
