## Supplementary figures and images for "A sample-to-identification workflow to monitor animal and plant biodiversity using metabarcoding and real-time analysis"

### RTBioScan_consensus_consolidated_tax_time.pdf

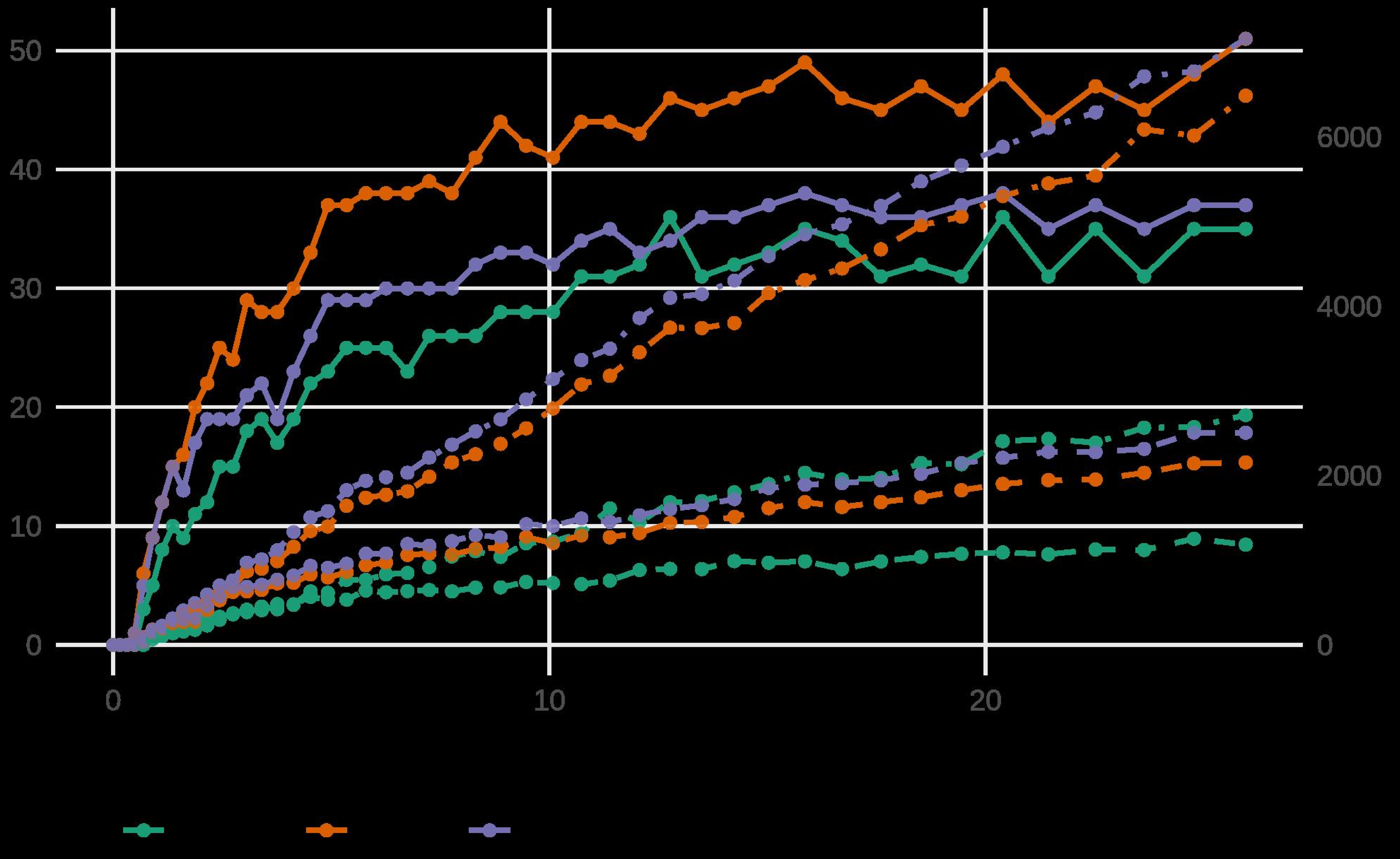

### RTBioScan_consensus_consolidated_tax_time.png

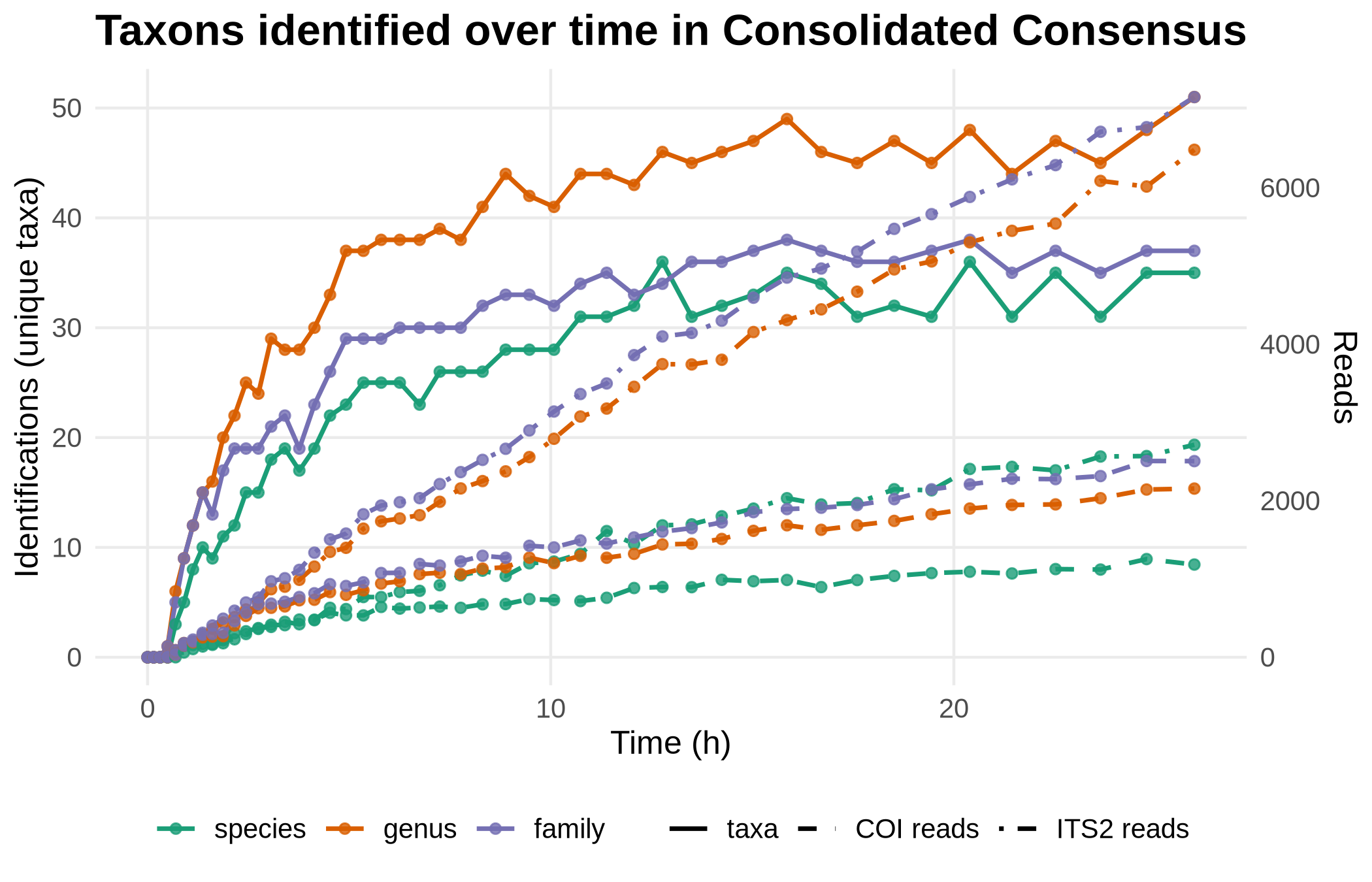

### RTBioScan_consensus_tax_time.pdf

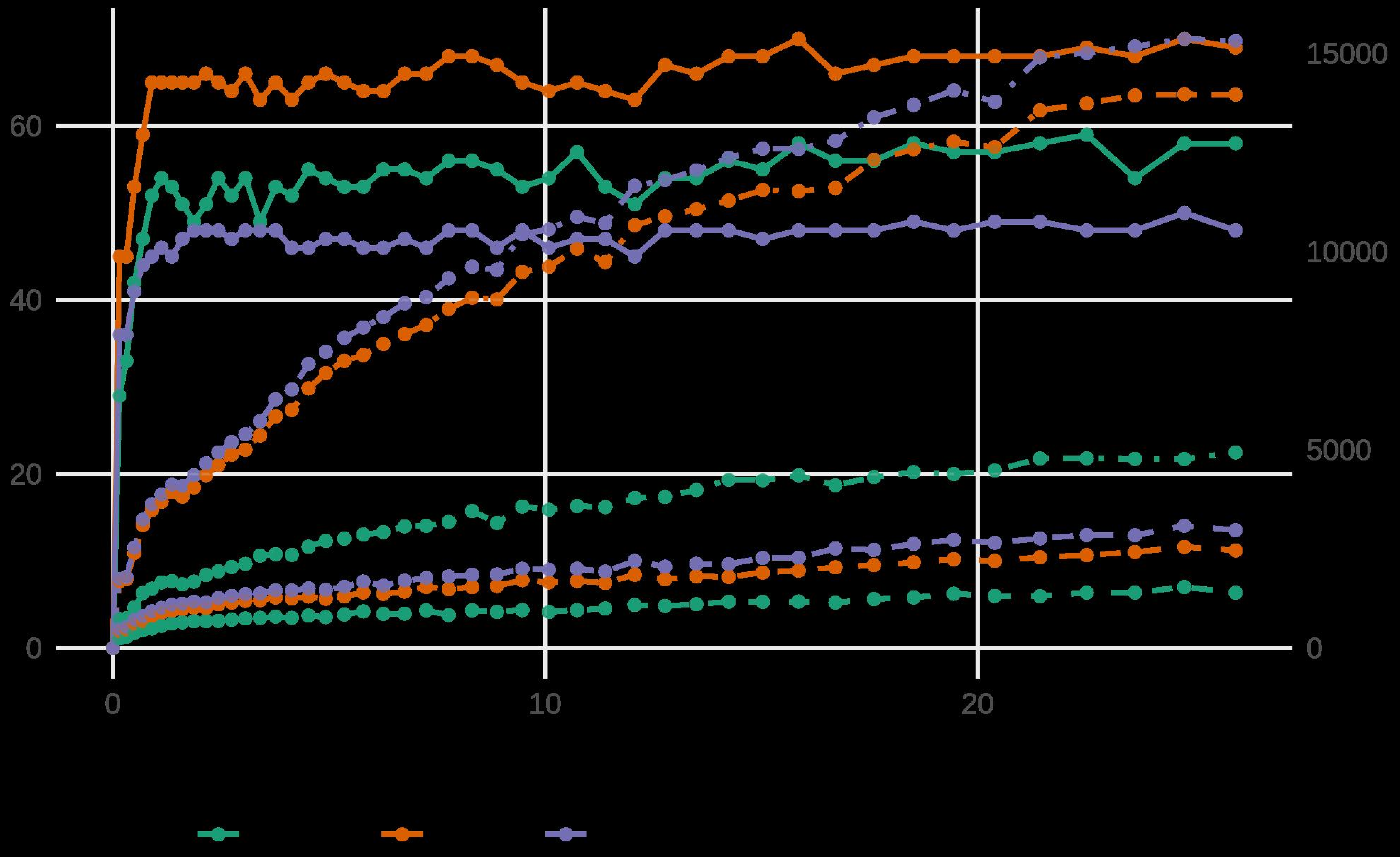

### RTBioScan_consensus_tax_time.png

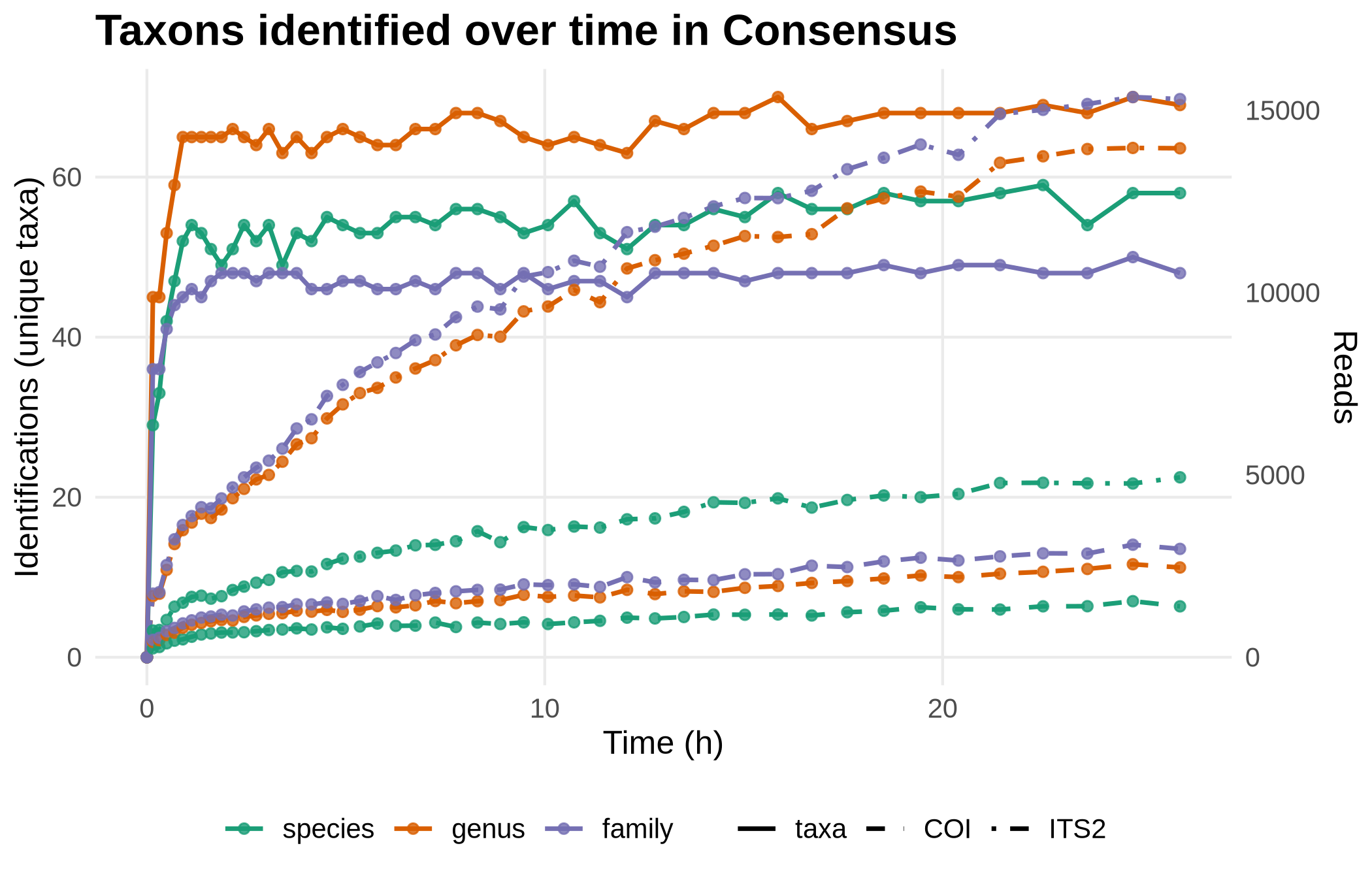

### RTBioScan_otu_frozen_tax_time.pdf

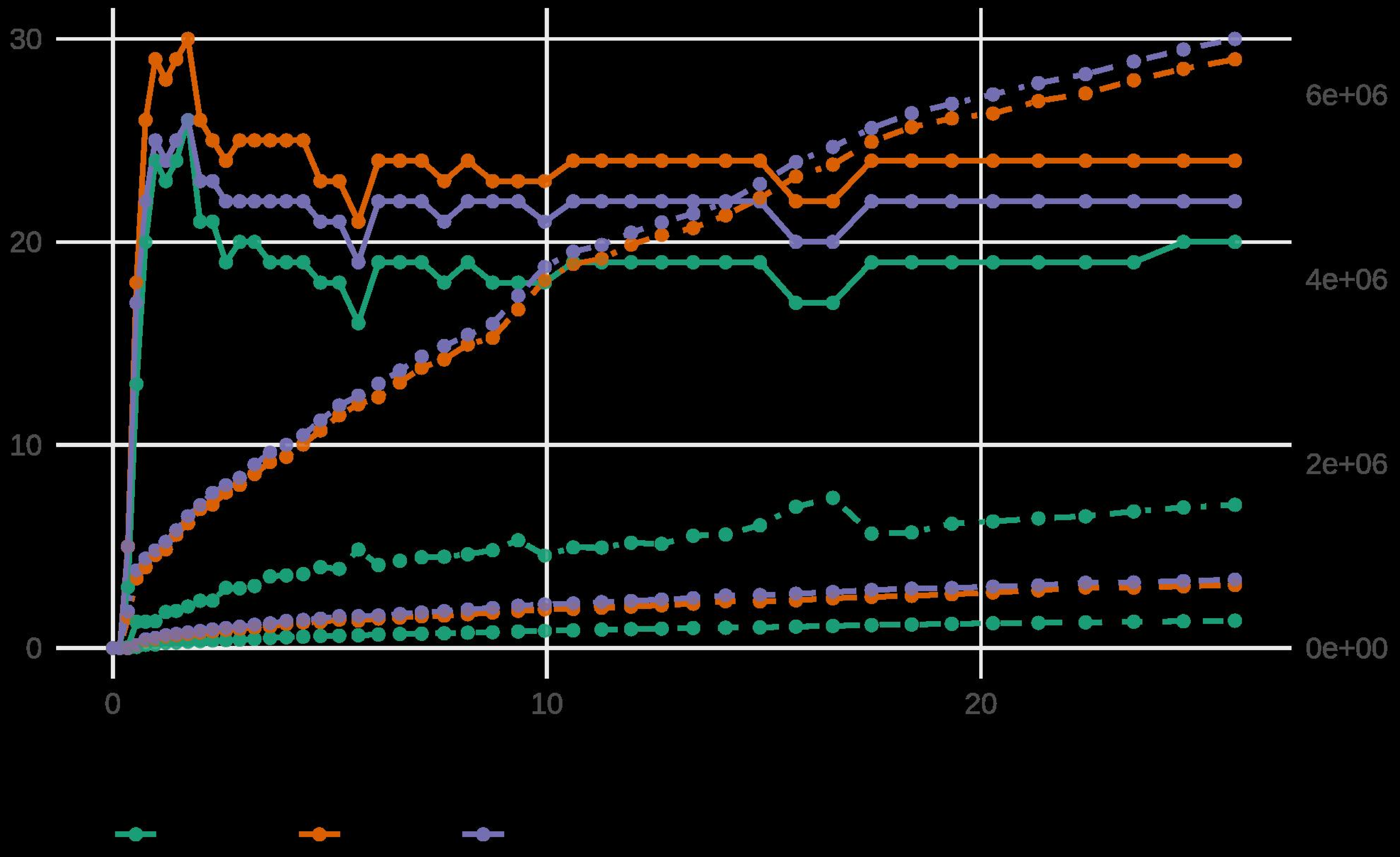

### RTBioScan_otu_frozen_tax_time.png

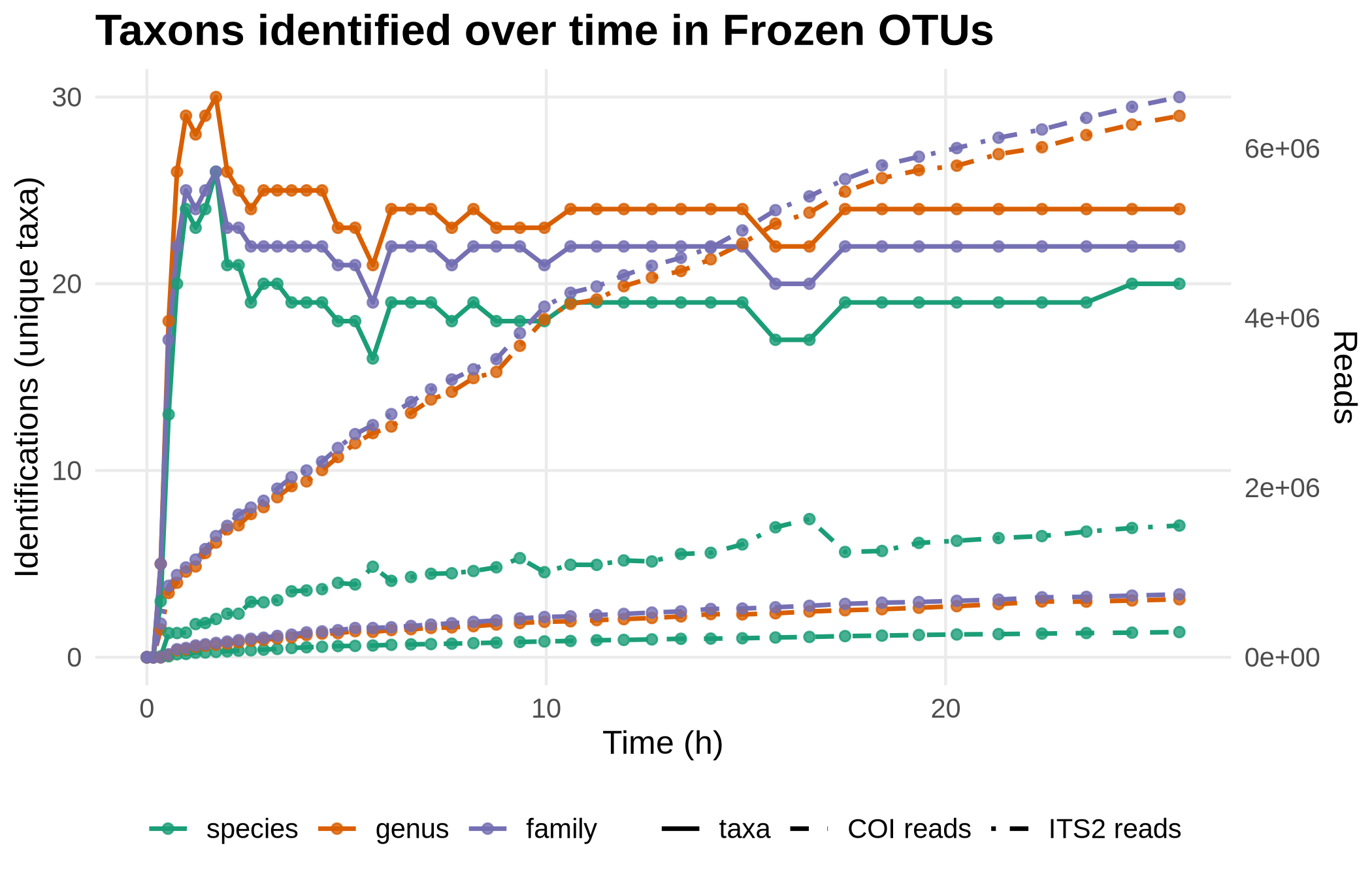

### RTBioScan_otu_tax_time.pdf

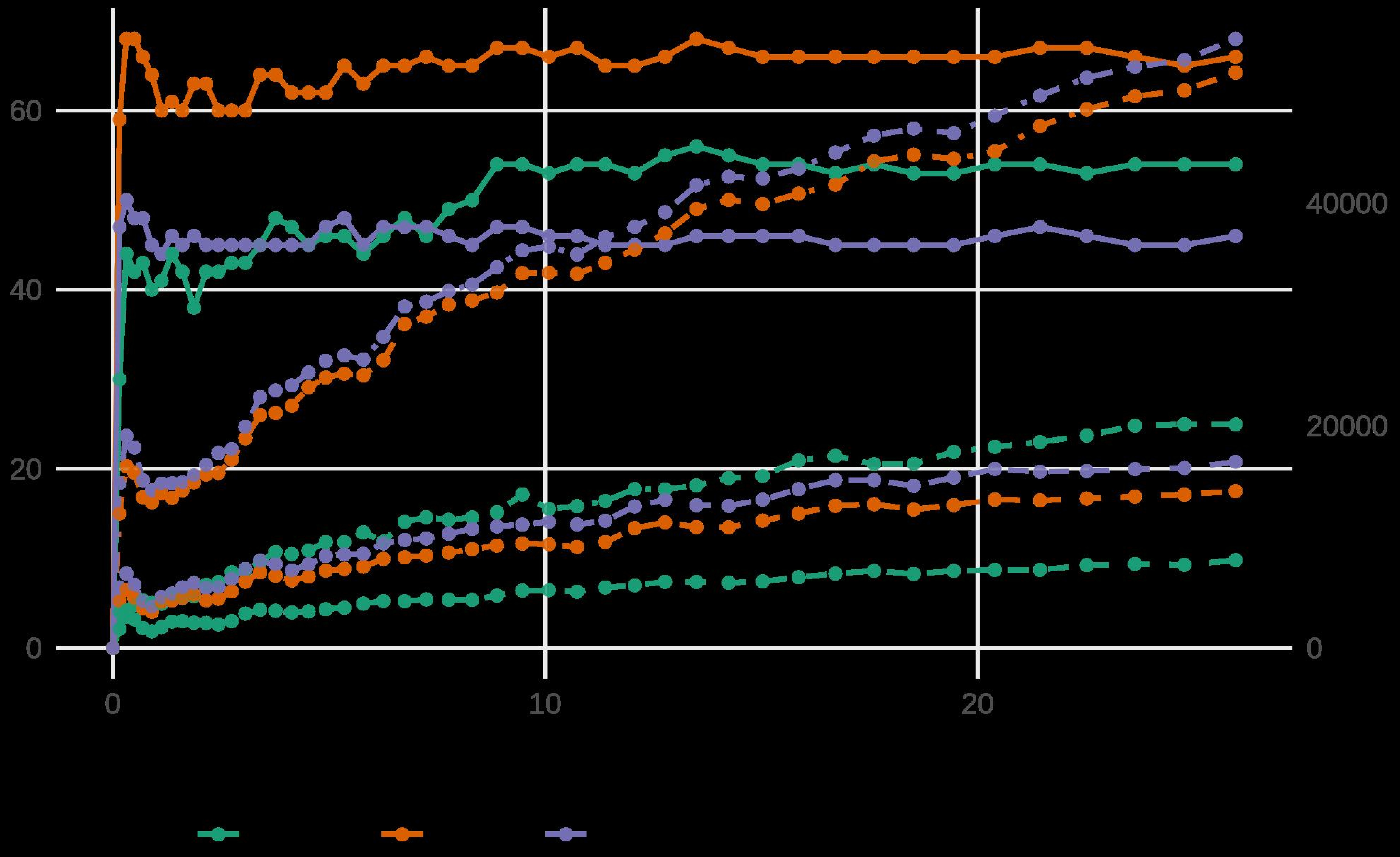

### RTBioScan_otu_tax_time.png

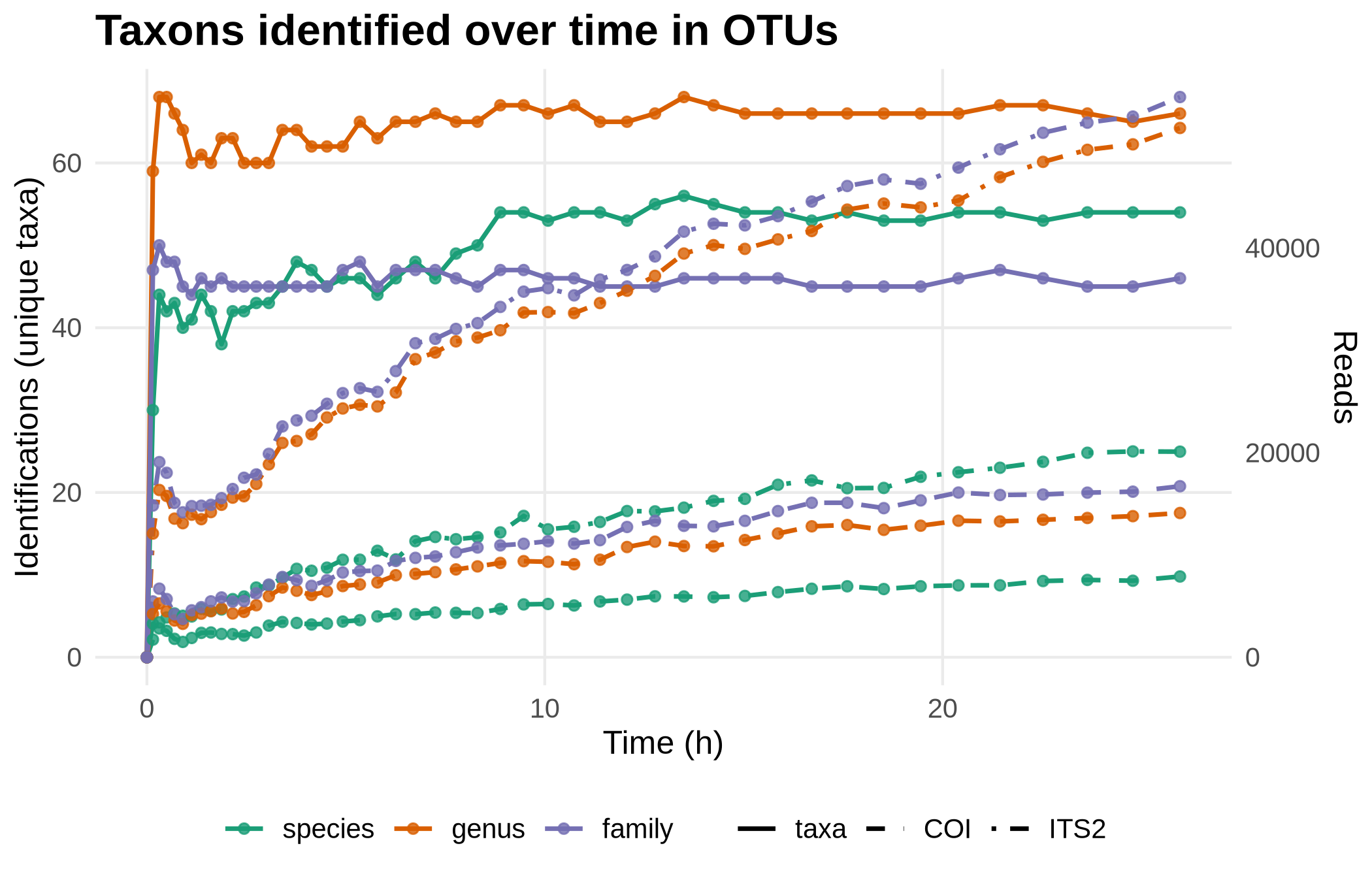

### RTBioScan_reads_cumulative_log.pdf

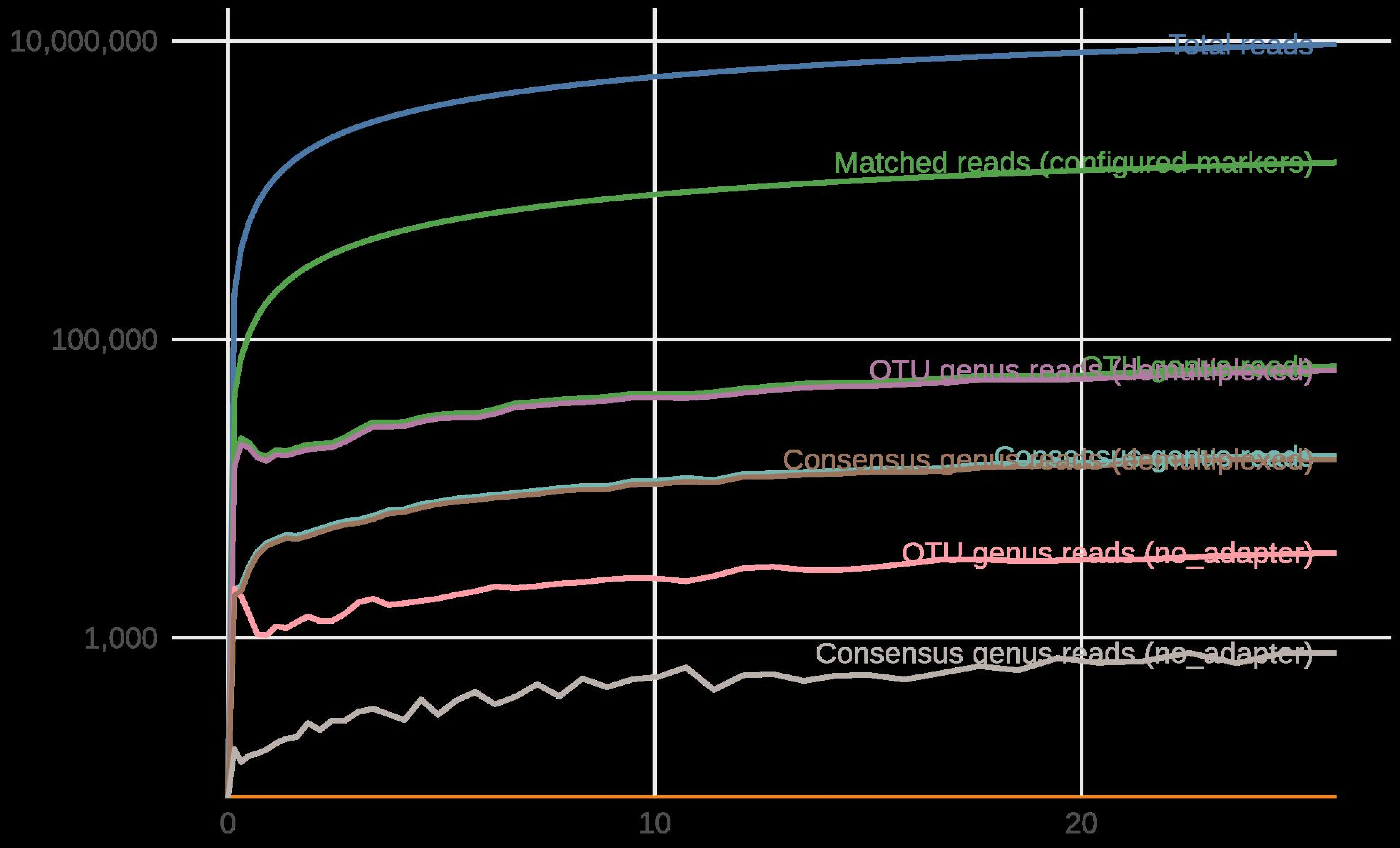

### RTBioScan_reads_cumulative_log.png

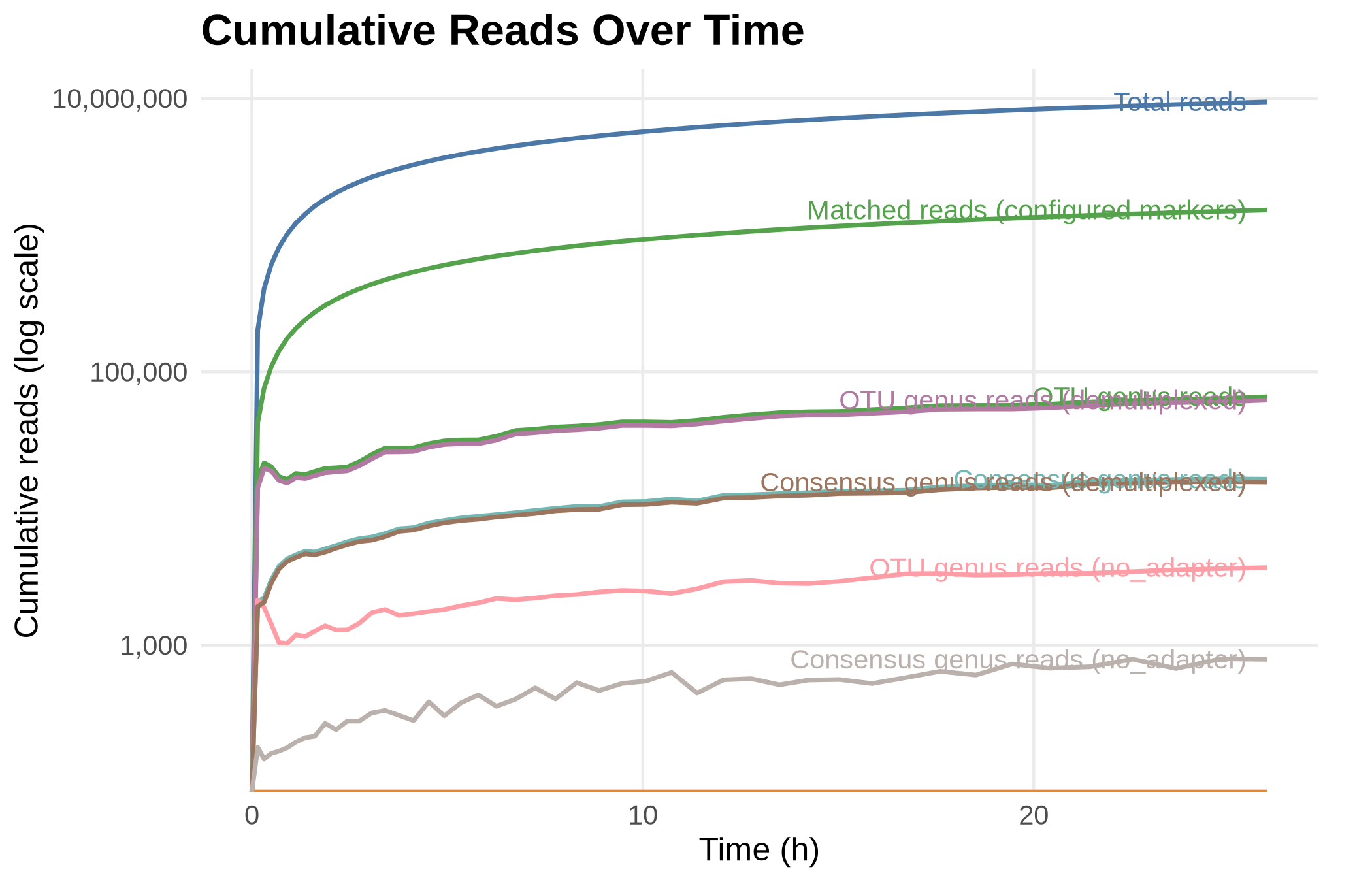

### RTBioScan_reads_time.pdf

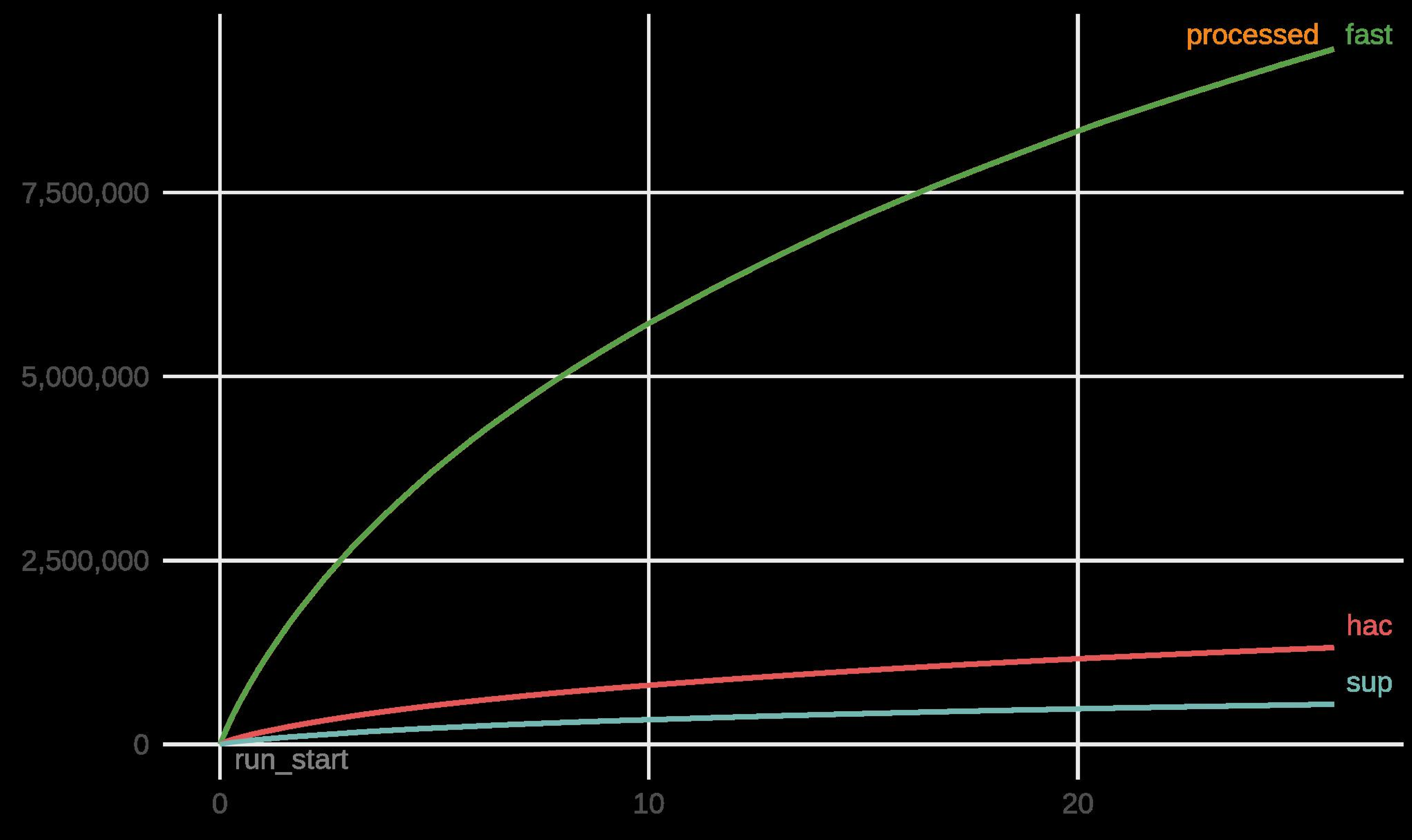

### RTBioScan_reads_time.png

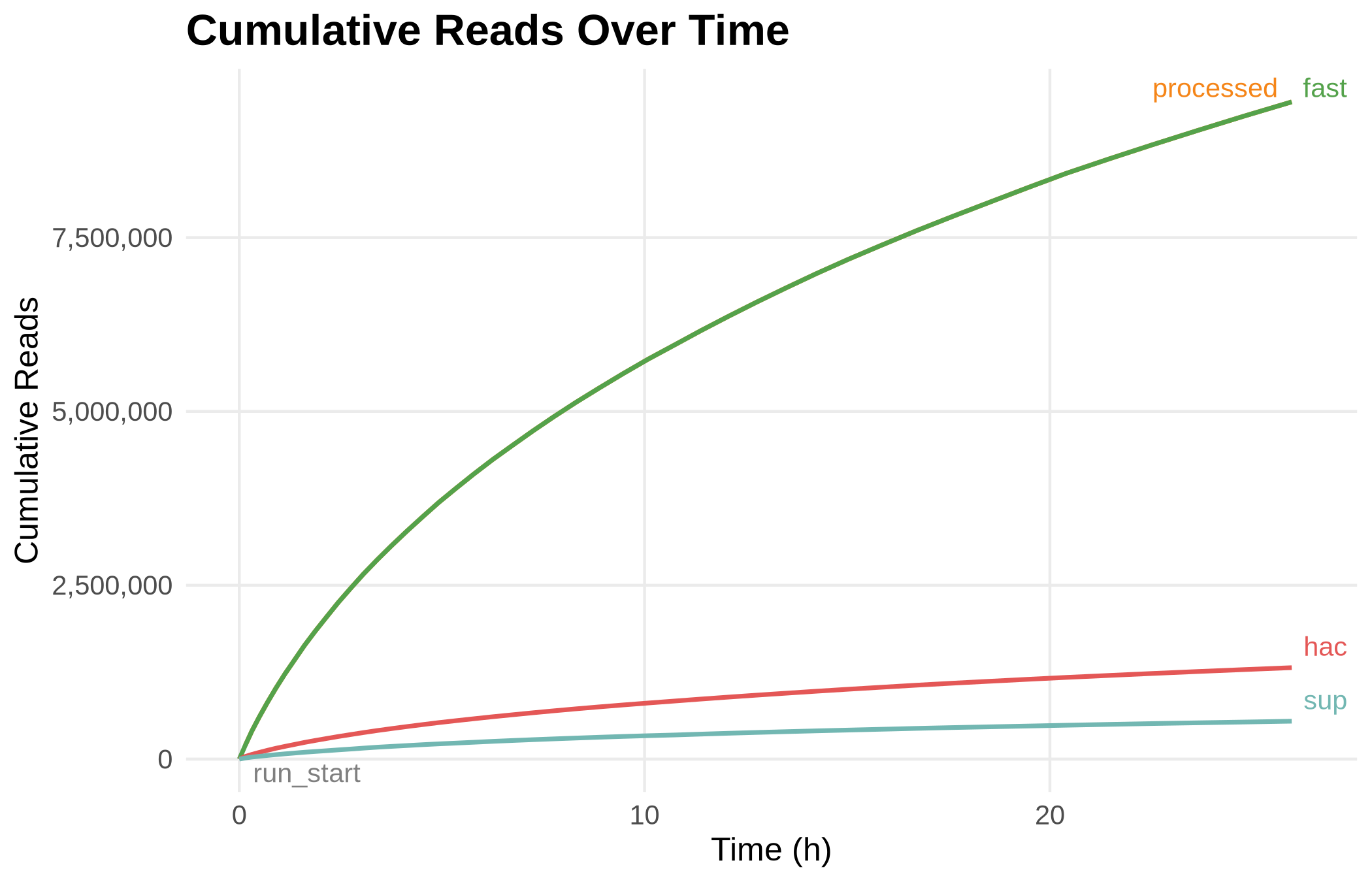
